# Discovery of tomato UDP-glucosyltransferases involved in bioactive jasmonate homeostasis using limited proteolysis-coupled mass spectrometry

**DOI:** 10.1101/2025.10.15.682356

**Authors:** Jhon Venegas-Molina, Lennart Mohnike, Sara Selma García, Hilde Janssens, Robin Colembie, Issl Kimpe, Ana Cristina Jaramillo-Madrid, Elia Lacchini, Johan M. Winne, Petra Van Damme, Ivo Feussner, Alain Goossens, Krešimir Šola

**Author notes:** These authors contributed equally (Alain Goossens, Krešimir Šola).

## Abstract

Jasmonic acid (JA) is the precursor of the bioactive molecule jasmonoyl-isoleucine (JA-Ile), a plant hormone that regulates fitness and development. Although JA biosynthesis, signaling, and responses have been intensively studied, the catabolism of JA remains incompletely understood. Here, we used the recently developed technique of limited proteolysis-coupled mass spectrometry (LiP-MS) to investigate metabolite–protein interactions in plants, aiming to discover enzymes involved in JA metabolism. We identified several previously reported JA-binding proteins, thus validating the robustness of the method, along with recognized enzymes of the JA pathway and a series of novel potential JA-binding proteins. We performed functional characterization of a set of identified JA-interacting UDP-glucuronosyltransferase (UGT) enzymes through omics, biochemical, enzymatic, and structural analyses. Our results demonstrate that two tomato UGTs effectively glucosylate JA to form JA-glucosyl esters, potentially playing a role in the regulation of bioactive JA homeostasis. With this, our findings uncovered a missing step in the metabolism of JA.

## Introduction

Jasmonic acid (JA) is the precursor of the bioactive molecule jasmonoyl-isoleucine (JA-Ile), a plant hormone that plays crucial roles in plant development and defense. In development JA regulates processes such as trichome formation, flower development, and root growth inhibition. In plant defense, for example, JA regulates the biosynthesis of specialized metabolites that serve as defensive toxins against necrotrophic pathogens and herbivorous insects^1,2^.

JA belongs to a group of compounds called oxylipins, which are metabolites derived from the oxidation of polyunsaturated fatty acids. The biosynthesis of JA begins in the plastids, where the fatty acid ɑ-linolenic acid is oxidized by 13-lipoxygenase to form 13-hydroperoxy-octadecatrienoic acid. This intermediate is then cyclized and rearranged by allene oxide synthase and allene oxide cyclase to form 12-oxo-phytodienoic acid (OPDA). OPDA is subsequently transported to the peroxisome, where it undergoes reduction by 12-OPDA reductase 3 and β-oxidation to ultimately form JA, which is then released into the cytosol^2^.

In the cytosol, JA undergoes metabolic conversions leading to various metabolites, each exhibiting distinct activities. On the one hand, there are few known bioactive JA-derivatives. For example, the conjugation of JA with isoleucine by the jasmonoyl amino acid conjugate synthetase JASMONATE RESISTANT 1 (JAR1) produces jasmonoyl-isoleucine (JA-Ile), considered the main and most efficient bioactive form of JA^3,4^. JA-Ile binds to the coreceptor complex consisting of the F-box protein CORONATINE-INSENSITIVE 1 (COI1) and repressor JASMONATE-ZIM DOMAIN (JAZ) proteins. JA-Ile sensing leads to the degradation of JAZ repressors, unleashing critical transcriptional regulators such as MYC2, which in turn drive the expression of genes that regulate development and defense^1,5–10^. JA can also be conjugated with alanine, leucine, methionine, and valine, each presenting different levels of bioactivity^11–13^. On the other hand, there are a handful of known inactive JA-derivatives. For example, JA is hydroxylated by JASMONATE-INDUCED OXYGENASEs (JOXs)^14^, to produce 12-OH-JA, which subsequently can undergo sulfation to form 12-HSO_4_-JA, or *O*-glucosylation to produce 12-*O*-Glucosyl-JA. These three JA-derivatives are inactive and tend to accumulate in wounded plants^15^. Bioactive JA-Ile can also be hydroxylated, for instance, by CYP94B3, to produce 12-OH-JA-Ile^16–18^. Other notable known JA-derivatives include methyl jasmonate (MeJA) and *cis*-jasmone. MeJA, an volatile metabolite produced in dicotyledonous plants by JA methyl transferases, is involved in JA communication to distal plant parts^19^; however, it becomes bioactive only after being cleaved by esterases and then converted into JA-Ile^20^. *Cis*-jasmone, the decarboxylated form of JA, is a volatile compound that accumulates following insect wounding and induces genes that produce volatile organic compounds used as defense mechanisms, such as attracting parasitoids against the attack of aphids^21^. Despite decades of intensive studies on JA metabolism, signaling, and responses, the number of JA-metabolizing enzymes identified to date remains limited. Consequently, our understanding of JA homeostasis, particularly JA catabolism, is still incomplete.

Limited proteolysis-coupled mass spectrometry (LiP-MS) is a recently developed method for detecting metabolite–protein interactions, initially applied in microbial and mammalian systems^22^. In LiP-MS, proteomes are extracted under native conditions and incubated with a metabolite of interest, followed by partial digestion using a nonspecific and promiscuous protease. Next, samples undergo shotgun sample preparation and are analyzed by mass spectrometry (MS). Finally, data analysis then identifies proteins containing differentially abundant peptides between samples incubated with the metabolite and those treated with buffer control. The presence of such peptides suggests that metabolite binding may protect specific protein regions from protease digestion or induce protein conformational changes, indicating potential metabolite–protein binding events^22^.

Here, we have adapted the LiP-MS methodology for use in plants and performed experiments with JA to discover potential JA-binding proteins in tomato (*Solanum lycopersicum*), and functionally characterized selected JA-binding proteins. Hereby, we demonstrate that two enzymes belonging to the UDP-dependent glycosyltransferase (UGT) family can glucosylate JA *in vitro* to form glucosyl esters. The activity likely contributes to an *in-planta* feedback loop to inactivate or store JA.

## Results

### LiP-MS identifies known and novel JA-binding proteins in tomato

In recent years, several large-scale proteomics techniques have emerged to identify and characterize protein–metabolite interactions^23,24^. Originally developed for microbial or mammalian proteomes, the implementation of these methods in plants often lags behind. This was also the case for LiP-MS, for which we have presented an adapted and accessible version for use in plants, as recently reported ^10^. Here, we exploited our plant LiP-MS platform to discover potential new JA-interacting proteins involved in JA metabolism or signaling in tomato. To this end, we extracted proteins from tomato hairy root cultures and performed LiP-MS with 1 and 10 mM JA. We identified 1,244 differentially abundant peptides in LiP-MS experiments with 10 mM JA, corresponding to 775 unique proteins. For experiments with 1 mM JA, we detected 183 differentially abundant peptides, corresponding to 154 proteins (Fig. 1). Of the latter set, 146 proteins (94.8%) were also found with 10 mM JA, demonstrating consistency across experiments conducted under related setups (Fig. 1).

**Figure 1.**
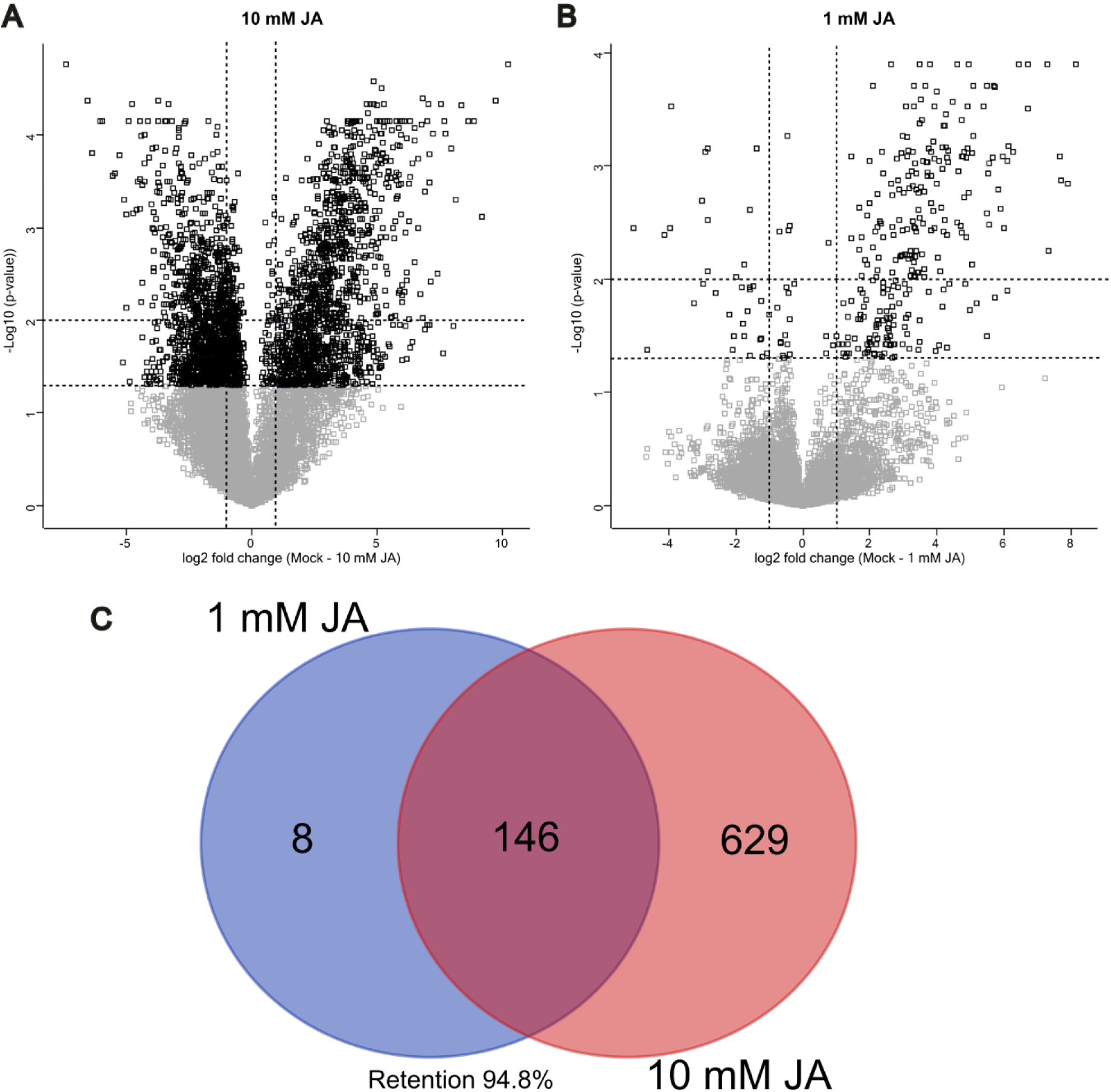
Output of LiP-MS with JA in tomato. A-B. Volcano plots depict log2-fold changes of peptide abundance as a function of statistical significance for 10 mM JA versus control (A) and 1 mM JA versus control (B). Differentially abundant peptides above significance cutoff values (adjusted p-value = 0.01 and 0.05; fold-change = 2) are highlighted in black (n = 4 biological replicates). (C) Venn diagram displaying the overlap of statistically significant proteins in samples treated with 10 mM JA versus 1 mM JA (FC > 2 and adjusted p-value < 0.01). Percentage retention is the percentage of protein interactors detected in both conditions.

Among the detected proteins with differential abundance (Supplementary Dataset 1), we identified the putative tomato homologs of a known Arabidopsis JA interacting protein, specifically several proteins with predicted peptidyl-prolyl cis-trans isomerase activity, the so-called cyclophilins. Using the reported Arabidopsis JA-binding Cyclophilin 20-3 (CYP20-3, At3g62030)^25^ as a reference for BLASTP searches, we identified its closest tomato homolog, protein A0A3Q7EA02 (Solyc01g009990), in our LiP-MS experiments as one of the peptidyl-prolyl cis-trans isomerases interacting with JA. Specifically, three peptides from this protein were differentially abundant in 10 mM JA LiP-MS setup (Supplementary Table 1). Throughout the manuscript, all mentions of peptides ‘identified by LiP-MS’ refer specifically to those that exhibit a statistically significant difference in abundance (e.g., q-value < 0.05) between experimental conditions, unless otherwise stated.

Regarding potential new JA-interacting proteins, we discovered proteins with known or predicted molecular functions that may be involved in JA metabolism. For instance, we identified a lipoxygenase (SlLOX, Solyc08g014000), a 12-oxophytodienoate reductase (SlOPR1, Solyc10g086220), an OPC-8:0-CoA ligase (SlOPCL, Solyc12g094520), an acyl-CoA oxidase (SlACX, Solyc10g008110), and a 3-ketoacyl CoA thiolase (SlKAT, Solyc09g061840) (Supplementary Table 2), all of which are enzymes known to be involved in JA biosynthesis^26^. Of note, this list also includes the enzymes (SlACX and SlKAT) of the final steps of the JA biosynthesis pathway, yielding JA as their enzymatic product. Though this might suggest possible hitherto unknown enzymatic feedback mechanisms, we were particularly intrigued by the numerous uridine diphosphate (UDP) glycosyltransferases (UGTs) detected. We selected these potential JA-interacting UGTs for deeper experimental characterization because, to our knowledge, glycosylation of JA has not been well characterized and may represent a novel JA catabolic pathway or storage mechanism. Specifically, in the setup using 10 mM JA, we identified peptides of twelve UGT proteins exhibiting significant differential abundance, four of which were also identified in the 1 mM JA setup (Table 1).

**Table 1.**
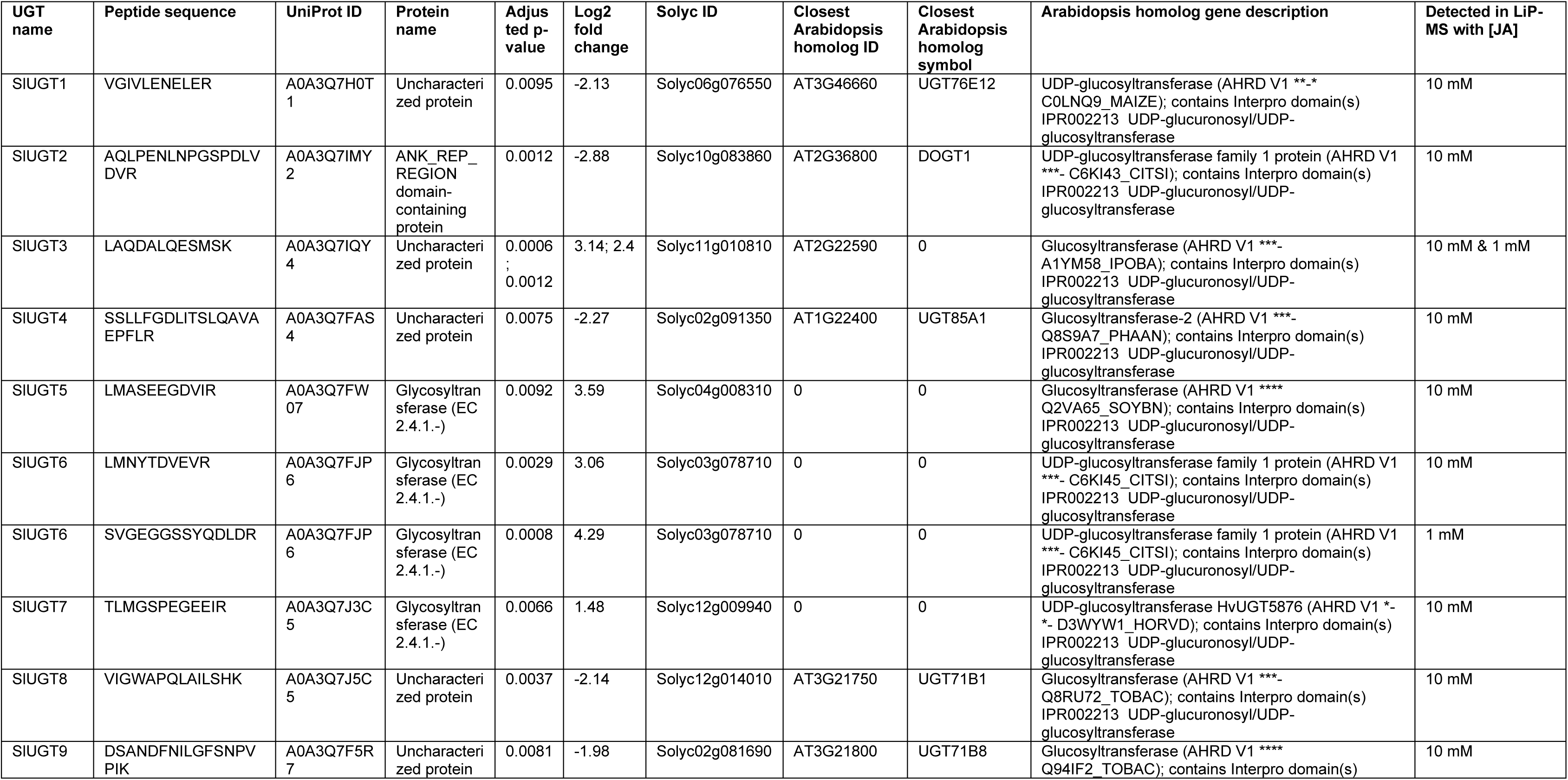
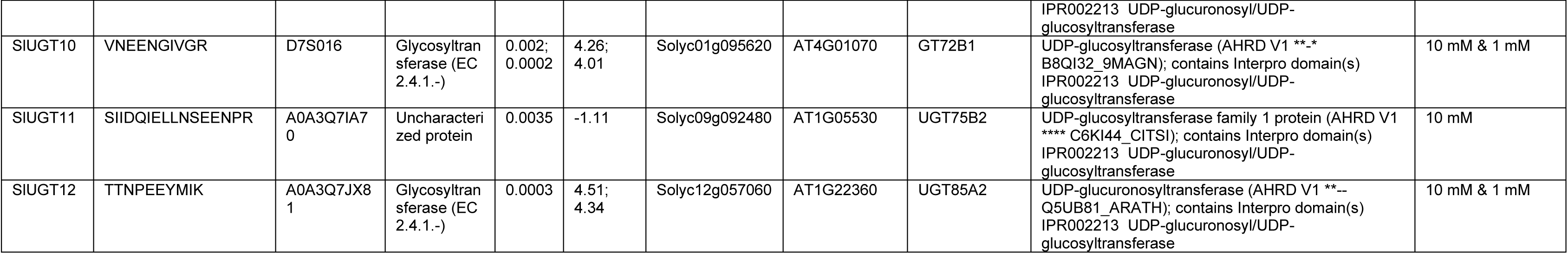
Peptides of putative interactor proteins detected with LiP-MS with JA with predicted UGT function. The table shows differentially abundant peptides and predicted functions based on their homology to Arabidopsis. If a peptide was detected in two experiments, it is listed in the final column, and its adjusted p-values and log2 fold changes from two treatments are separated by a semicolon.

To narrow down the list of potential JA-glycosylating UGTs, we assessed whether the genes corresponding to the twelve UGTs identified by LiP-MS were transcriptionally regulated upon JA induction. The rationale behind this selection criterion is that the expression of genes encoding enzymes involved in JA metabolism is typically transcriptionally controlled by JA itself, including both biosynthesis and catabolism, as part of amplification and feedback loops, respectively^5,6,9,27^. Therefore, we consulted in-house generated RNA-Seq data from tomato hairy root cultures treated with 50 µM JA and sampled at three timepoints over a 24-hour period (30 min, 4 h and 24 h)^28^. Notably, for five out of the twelve UGTs identified by LiP-MS, JA-modulated expression was observed. More specifically, the expression of *SlUGT8, SlUGT9, SlUGT10, SlUGT11*, and *SlUGT12*, was upregulated by JA treatment at the three timepoints sampled (with fold-change > 2 and FDR < 0.05, Supplementary Dataset 2) (Fig. 2). The transcript *Solyc11g010810* (*SlUGT3*) was not detected in any sample in our RNA-Seq dataset, whereas the other six verified *UGT* genes showed no response to JA (Fig. 2). Since *SlUGT8, SlUGT9, SlUGT10, SlUGT11*, and *SlUGT12* were rapidly induced by JA, we hypothesized that these UGTs are more likely to be involved in JA metabolism, more particularly in a feedback loop to dampen the JA signal. Based on this reasoning, we decided to focus further on those five JA-inducible UGTs.

**Figure 2.**
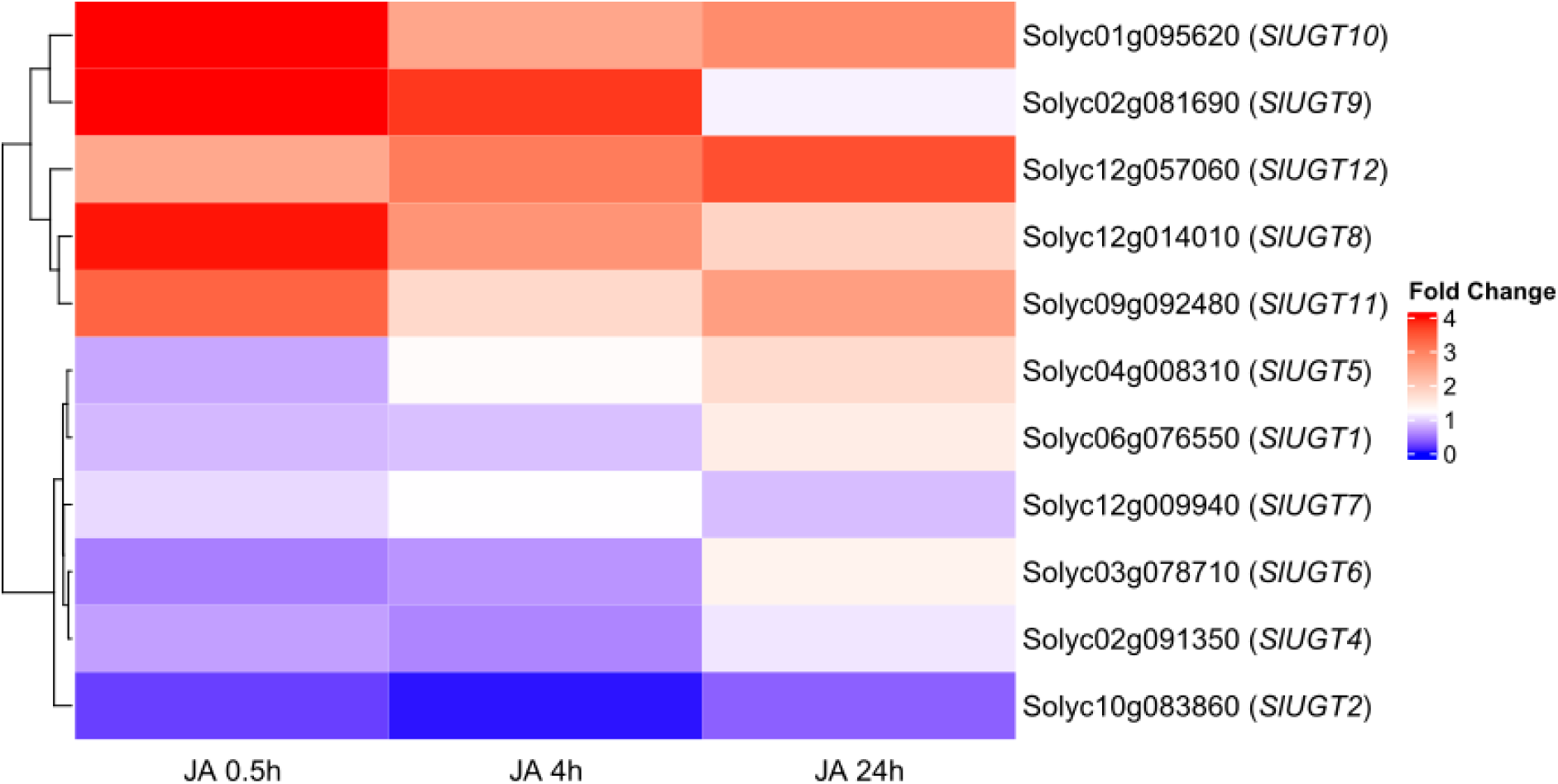
Genes encoding potential JA-binding UGTs are transcriptionally regulated by JA in tomato roots. Heatmap depicting gene expression patterns of potential JA-binding UGTs detected with LiP-MS. Two-week-old tomato hairy root cultures were treated with 50 µM JA or mock, and samples were collected after 0.5, 4, or 24 h. The heatmap depicts the fold-change of JA treated samples compared to mock for 12 selected tomato UGTs. Upregulation is depicted in red (fold-change > 2 and FDR < 0.05), and no effect in blue (n = 3).

### Correlations between peptides detected by LiP-MS and predicted JA-binding sites in JA-inducible UGTs

First, we sought to structurally analyze the five selected UGTs *in silico*. Specifically, we aimed to determine whether the locations of the peptides detected by LiP-MS within the UGT proteins correspond to potential JA-binding sites. Additionally, we examined whether the structural features of the five JA–UGT interactions aligned with known features of putative catalytic domains.

We retrieved the predicted UGT protein structures using AlphaFold^29^ and obtained the 3D structure of JA from PubChem^30^. JA–UGT interaction models were constructed via molecular docking using COACH-D^31^. Each metabolite–protein structure model generated a confidence score (C-score) ranging from 0 to 1, indicating the reliability of the predictions. Using these structural models, we then compared the positions of the peptides detected by LiP-MS with the JA–UGT binding sites predicted by COACH-D. To obtain a high sequence coverage, we included all peptides identified in the LiP-MS experiment with 10 mM JA for the five selected UGTs (Supplementary Table 3) but highlighted only those peptides that passed the statistical threshold of q-value < 0.05 (Fig. 3; Supplementary Fig. 1-4). Lastly, we measured the atomic distances between JA and the peptides detected by LiP-MS, comparing them to distances of established catalytic interactions. According to calculations by Piazza, et al. ^22^, atomic distances of 6.44 Å from the metabolite can define boundaries of active sites, thereby suggesting a catalytic nature of the metabolite–protein interactions observed in LiP-MS experiments.

**Figure 3.**
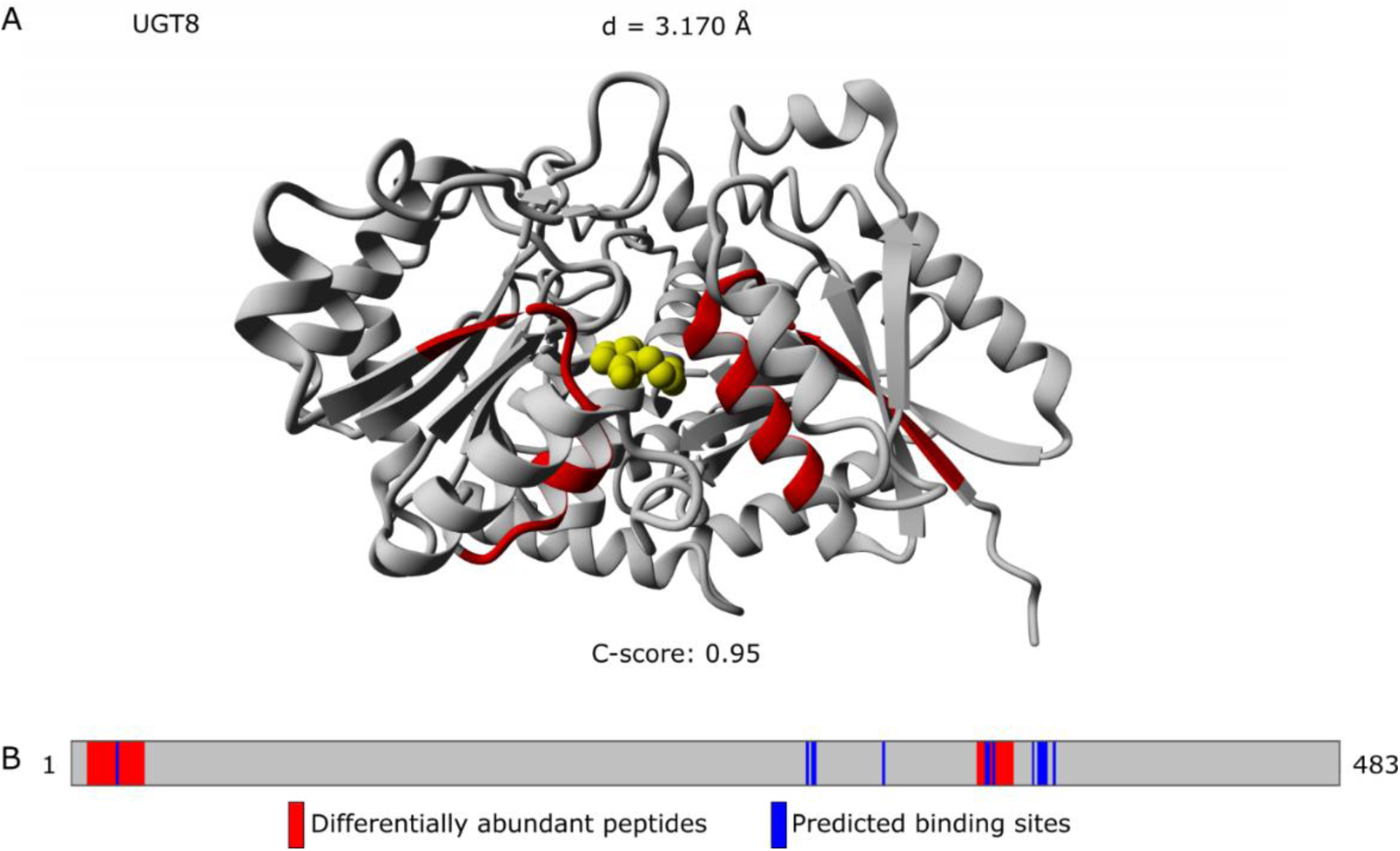
3D metabolite–protein interaction structural model of JA–SlUGT8. A. 3D structural model showing the interaction between JA (yellow) and SlUGT8 (gray), with differentially abundant peptides detected by LiP-MS highlighted in red. B. Protein map illustrating the overlap between the differentially abundant peptides detected by LiP-MS (red) and the COACH-D predicted binding sites (blue). C-score: COACH-D model confidence score as defined by Wu *et al*.^31^. d: minimal distance between the atoms of JA and the differentially abundant peptides detected by LiP-MS.

Eighty percent of the 3D structure models built with COACH-D had high-confidence prediction scores. In most cases, the peptides detected by LiP-MS and the JA-binding sites predicted by COACH-D were located within protein regions corresponding to the putative substrate-binding site (Supplementary Fig. 2-4). Differentially abundant peptides from SlUGT8, SlUGT9, and SlUGT10 overlapped with the predicted JA-binding sites (Fig. 3; Supplementary Fig. 1-2), demonstrating a correlation between the LiP-MS data and the *in silico* metabolite–protein interaction predictions. Additionally, the minimal atomic distances between JA and peptides detected with LiP-MS for SlUGT8, SlUGT10, and SlUGT11 suggest potential catalytic-type interactions (Fig. 3; Supplementary Fig. 2-3). These results provide strong evidence that the interactions between JA and the UGTs identified through LiP-MS are likely to be functionally relevant, potentially playing a role in JA metabolism.

### SlUGT8 and SlUGT11 use JA as a substrate to form jasmonyl-1-β-glucose (JA-Glc) *in vitro*

To assess enzymatic activity, we recombinantly expressed the SlUGT proteins. Therefore, each SlUGT was fused to a C-terminal 6x His-tag and heterologously expressed in *Escherichia coli.* We successfully expressed and purified SlUGT8, SlUGT10, SlUGT11, and SlUGT12 (Supplementary Fig. 5-8). In contrast, recombinant SlUGT9 showed very low to no expression, and its presence was observed only in insoluble fractions. Hence, SlUGT9 was excluded from further experimental characterization.

Next, we performed enzyme activity assays with the purified recombinant UGTs to test their ability to glycosylate JA. When SlUGT8 and SlUGT11 were incubated with JA and UDP-glucose (UDP-Glc) overnight and then the reaction mixtures were subjected to LC-MS analysis, a new chromatographic peak distinct from the negative control appeared (Fig. 4A). This peak was inferred to be corresponding to JA-Glc based on the mass of precursor and product ions (Fig. 4C). No differential peaks were detected for SlUGT10 and SlUGT12, suggesting that these heterologously expressed enzymes are unable to glucosylate JA, at least under the *in vitro* conditions tested. SlUGT8 catalyzed the synthesis of JA-Glc within 30 min (Supplementary Fig. 9), suggesting that it may be more efficient at catalyzing the reaction than SlUGT11 when heterologously expressed. Additionally, unlike SlUGT11, SlUGT8 was able to glucosylate hydroxylated JA (12-OH-JA) to form 12-OH-JA-Glc (Fig. 4B and D). Neither SlUGT10 nor SlUGT12 glycosylated either of the tested substrates.

**Figure 4.**
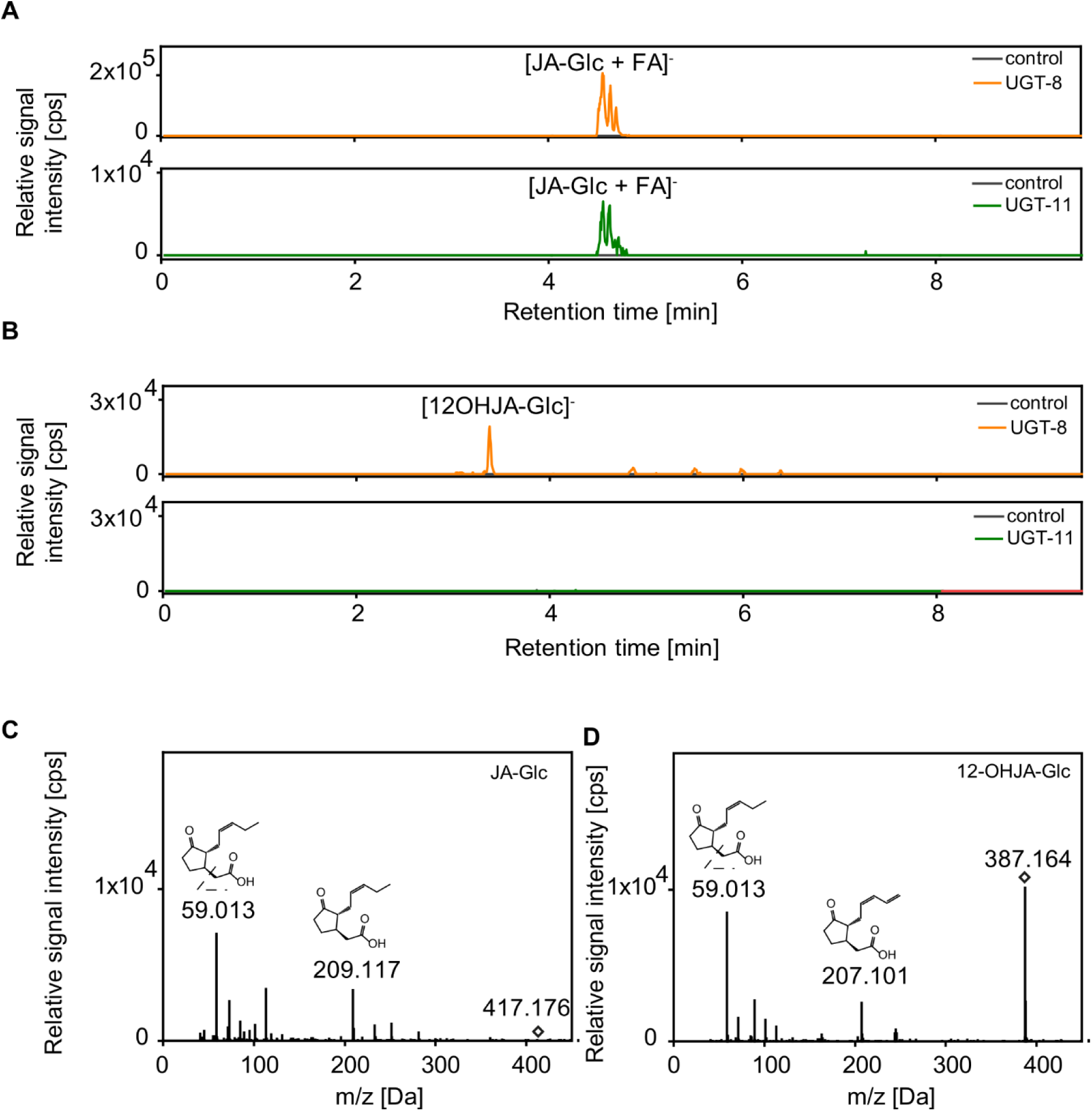
LC-MS-based activity assays of SlUGTs with JA and 12-OH-JA. A. Extracted ion chromatograms of JA activity assay using 0.5 mM JA and 0.5 mM UDP-Glc as substrates for purified SlUGT8 and SlUGT11. B. Extracted ion chromatograms of 12-OH-JA activity tests using 0.5 mM 12-OH-JA and 0.5 mM UDP-Glc as substrates for purified SlUGT8 and SlUGT11. C. JA-Glc precursor and product ions resulting from activity assays. D. 12-OH-JA-Glc precursor and product ions resulting from activity assays. Extracted ion chromatograms are shown for the product JA-Glc and 12-OH-JA-Glc. Signal intensity is given in counts per second (cps).

We further validated the activity SlUGT8 activity using the UDP-Glo™ Glycosyltransferase Assay kit (Promega), which employs bioluminescence as a readout of glycosylation activity. SlUGT8 activity increased with increasing enzyme concentration (Supplementary Fig. 10), indicating a direct dependence of catalysis on SlUGT8. Further, SlUGT8 used UDP-Glc as a sugar donor but not UDP-glucuronic acid (UDP-GlcA). These results suggest that UDP-Glc is the substrate for SlUGT8, while UDP-GlcA is not.

### Exogenous JA application and wounding trigger the accumulation of glucosyl esters of JA in tomato

To test whether the glycosylation of JA is a genuine part of JA homeostasis metabolism in tomato, we conducted untargeted metabolomics on tomato hairy root samples treated with exogenous JA. We identified the accumulation of two glycosylated JA-derivatives, namely JA-hexose (JA-Hex) and JA-Hex-Hex (data not shown), both of which are potential enzymatic products of the UGTs identified by LiP-MS. We then performed targeted metabolite analysis to quantify these compounds, using an in-house synthesized glucosyl ester of JA (JA-Glc) as a standard (Supplementary Fig. 11). For this experiment, tomato hairy root liquid cultures were treated with 10 μM JA, 50 μM JA, or mock (ethanol) for 3, 6, 12, and 24 h. Extracted metabolites were analyzed using LC-Q-TOF-MS/MS. The relative abundances of JA-Hex (confirmed as JA-Glc) and JA-Hex-Hex were compared across treatments and expressed as detector response per mg fresh weight (relative abundance). JA-Glc and JA-Hex-Hex exhibited different relative abundances across treatments and time points (Fig. 5). Both compounds were undetectable in mock-treated roots but accumulated markedly following JA treatment, suggesting they are derived directly from exogenous JA (possibly appended with *de novo* produced JA following the amplification loop^5,6,9,27^). *UGT* genes, such as *SlUGT8* and *SlUGT11*, encoding the enzymes responsible for this conversion, were also transcriptionally induced under these conditions.

**Figure 5.**
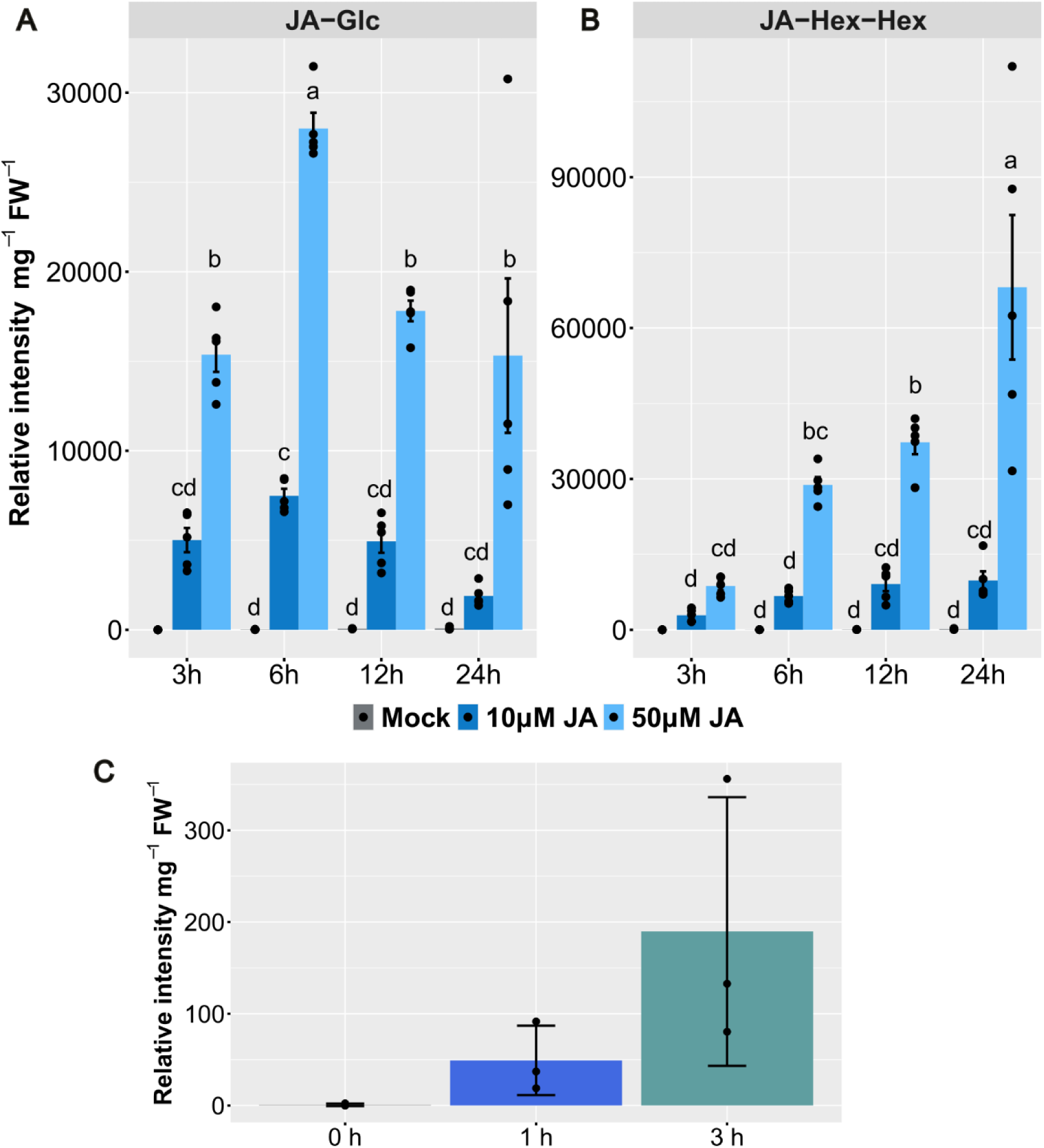
Relative quantities of jasmonyl-1-β-D-glucose (JA-Glc) and jasmonyl-1-dihexose (JA-Hex-Hex) in tomato hairy roots treated with JA and wounded tomato leaves. A-B. Levels of JA-Glc (A) and JA-Hex-Hex (B) in two-week-old tomato hairy roots treated with mock, 10 µM JA, or 50 µM JA, and harvested 3, 6, 12, or 24 h after treatment (n = 5). Lower case letters represent statistical differences between treatments according to Tukey’s HSD test, which was done following one-way ANOVA (p-value < 0.05). C. Levels of JA-Glc in leaves of 5-week-old tomato plants 0 h, 1 h and 3 h after wounding (n = 3).

Levels of JA glucosyl esters increased with higher concentrations of JA and were synthesized within 3 h. JA-Glc levels plateaued after 6 h (Fig. 5A), suggesting that JA-Glc may either be converted into another metabolite or degraded over time. Unlike JA-Glc, JA-Hex-Hex levels continued to rise throughout the treatment period (Fig. 5B). Since it is unlikely that JA-Glc would degrade while JA-Hex-Hex remains stable, it is plausible that JA-Hex-Hex is synthesized from JA-Glc, making it a JA-Glc-Hex compound. This is consistent with the observation that both JA-Glc and JA-gentiobiose (Ja-Glc-Glc) occur in tobacco BY-2 cells treated with JA^32^.

To ensure that the accumulation of JA-Glc is not an artefact of the exogenous JA used in treatments, we tested whether plants accumulate JA-Glc upon wounding. Whereas at the time of wounding plants did not have detectable JA-Glc, it appeared over time, with 3 h timepoint having more JA-Glc than 1 h timepoint (Fig. 5C; Supplementary Fig. 12), indicating that JA-Glc accumulates as wounding-induced JA accumulates. This shows that JA-Glc can naturally occur in tomato plants.

### JA-Glc has less effect on root growth and transcriptional activation of JA-responsive genes than JA

The inhibition of root growth was one of the first discovered physiological effects of exogenous JA application and has since been widely used as a characteristic trait in mutant screens and bioassays^33–36^. Given the endogenous JA-glucosylating activity observed with exogenously applied JA, we hypothesized that glucosylation might be one mechanism to inactivate JA. To test this, we chemically synthesized JA-Glc and treated tomato seedlings with JA-Glc or JA to compare their effects on root growth inhibition. After germination, seedlings were transferred to control media, or media containing 1 µM JA, or 1 µM JA-Glc. Both after two days (Fig. 6A and B) and three days (Fig. 6C and D), differences between treatments were observed. As expected, JA significantly inhibited root growth compared to the control treatment. Notably however, the effect of JA-Glc was less pronounced (Fig. 6B-D). This indicates that JA-Glc is less effective than JA at root growth inhibition.

**Figure 6.**
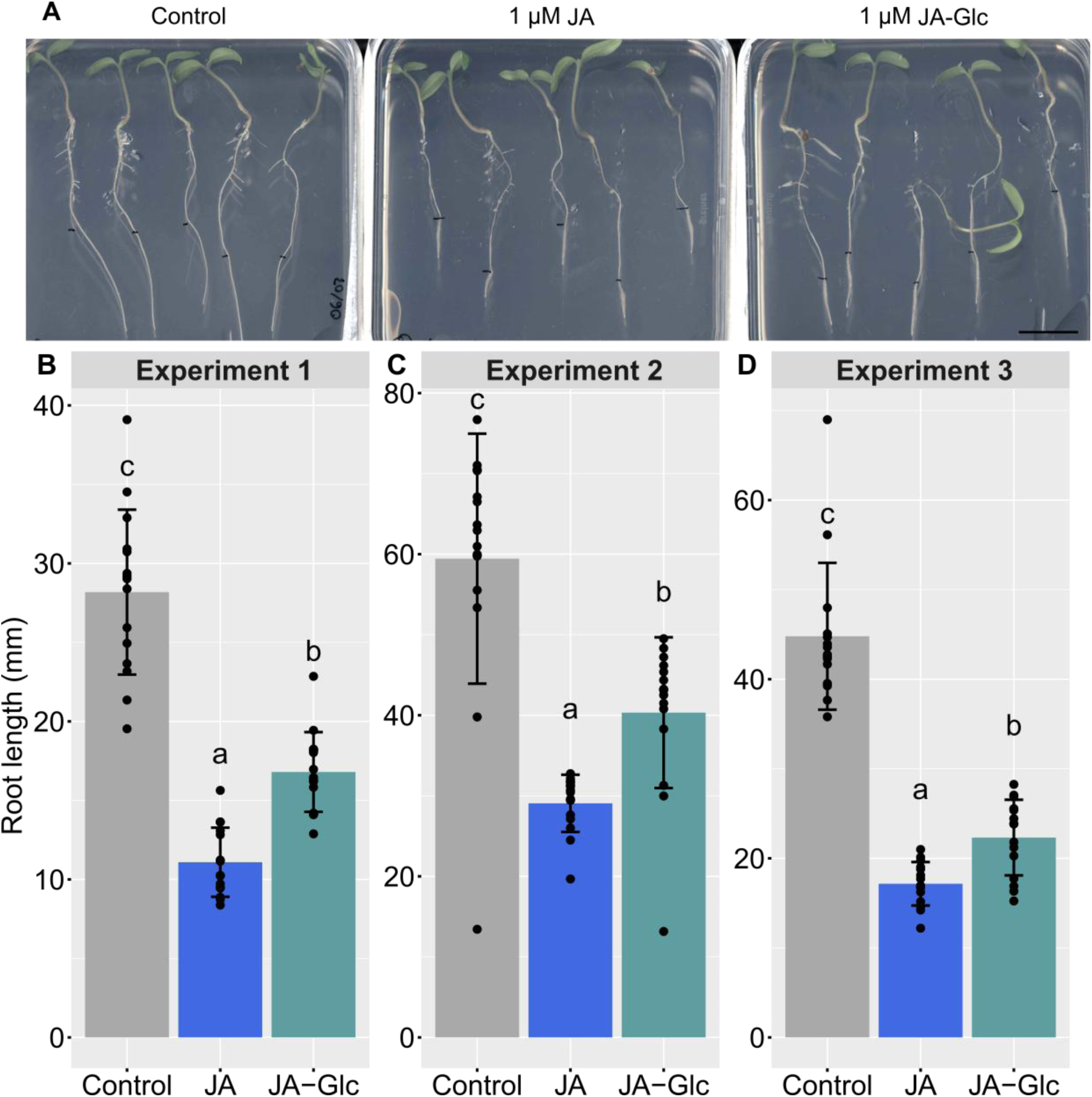
Root growth bioassay with application of JA or JA-Glc in tomato. A-D. One-week-old tomato seedlings were transferred to MS medium supplemented with control, 1 µM JA, or 1 µM JA-Glc. Root length was measured after two (A and B) or three (C and D) days of treatment. A. Images illustrating root growth under control, 1 µM JA, and 1 µM JA-Glc treatments. Black marks in each root indicate the initial root length at the time of metabolite application. Scale bar = 2 cm. B-D. Bar plots depicting root length after two (B) or three (C and D) days of control, 1 µM JA, and 1 µM JA-Glc treatments. Black dots represent values for individual roots, bars represent the means, and error bars the standard deviation. Letters above bars display groupings according to Tukey’s HSD test for multiple comparisons that was performed after one-way ANOVA (n = 15; p-value < 0.05).

To assess whether JA-Glc is also less effective at activating JA-responsive genes than JA, we measured transcript levels of three JA-inducible genes (*JAZ1*, *JAZ6*, and *JAZ7*) after a 6-h treatment with control, 1 µM JA, or 1 µM JA-Glc using qRT-PCR (Fig. 7, Supplementary Fig. 13). All three genes were induced by JA-Glc to lower levels than by JA, demonstrating that glycosylation of JA reduces its signaling activity. In one experiment (Supplementary Fig. 13), no statistically significant difference was observed between JA and JA-Glc treatment for *JAZ1*. This is likely due to high variability between replicates in the JA treatment group, which however, did not affect the results for the much more responsive *JAZ7*.

**Figure 7.**
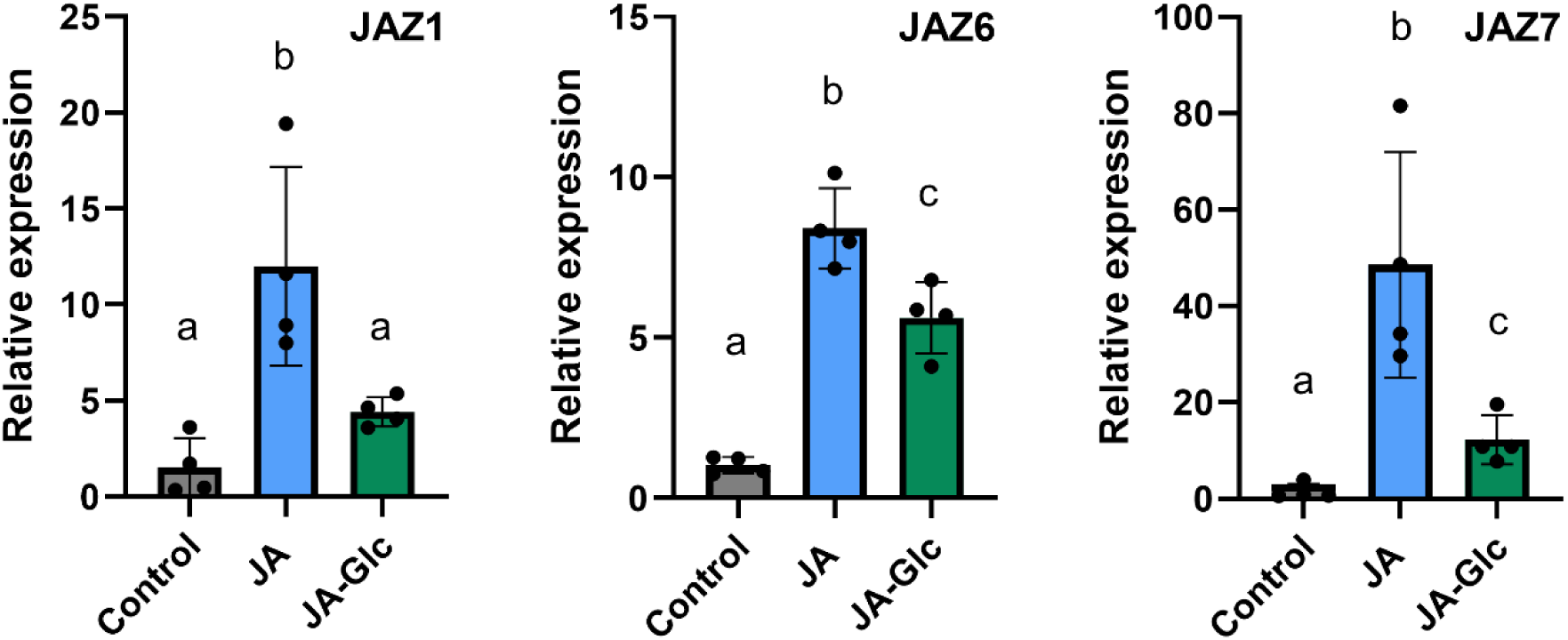
qRT-PCR analysis of the JA-responsive marker genes in tomato roots after treatment with JA and JA-Glc. Four plates with eight wild-type roots pooled per plate (biological replicates; n = 4) were employed for each 6-h treatment with 1 µM JA, 1 µM JA-Glc, or ethanol as control. Black dots represent values for each biological replicate, bars represent the means, and error bars the standard deviation. Lower case letters represent statistical differences between treatments calculated by Tukey’s HSD test, following one-way ANOVA (p-value < 0.05).

Based on these findings, we postulate that glycosylation of JA either inactivates it or at least partially reduces its biological activity.

### Generation of loss-of-function tomato hairy root lines to study the *in-planta* **roles of *UGT*s in JA responses**

To better understand the biological functions of the selected UGTs, we generated tomato hairy root lines with mutations in *UGT*s using CRISPR/Cas9. To generate the CRISPR/Cas9-based *UGT* loss-of-function tomato lines, we designed two guide RNAs (gRNAs) targeting sequence regions within the first exon of each *SlUGT* (Fig. 8A) following the protocol by Swinnen, et al. ^37^. CRISPR mutagenesis was performed targeting each single gene, as well as multiplexing all five at once.

**Figure 8.**
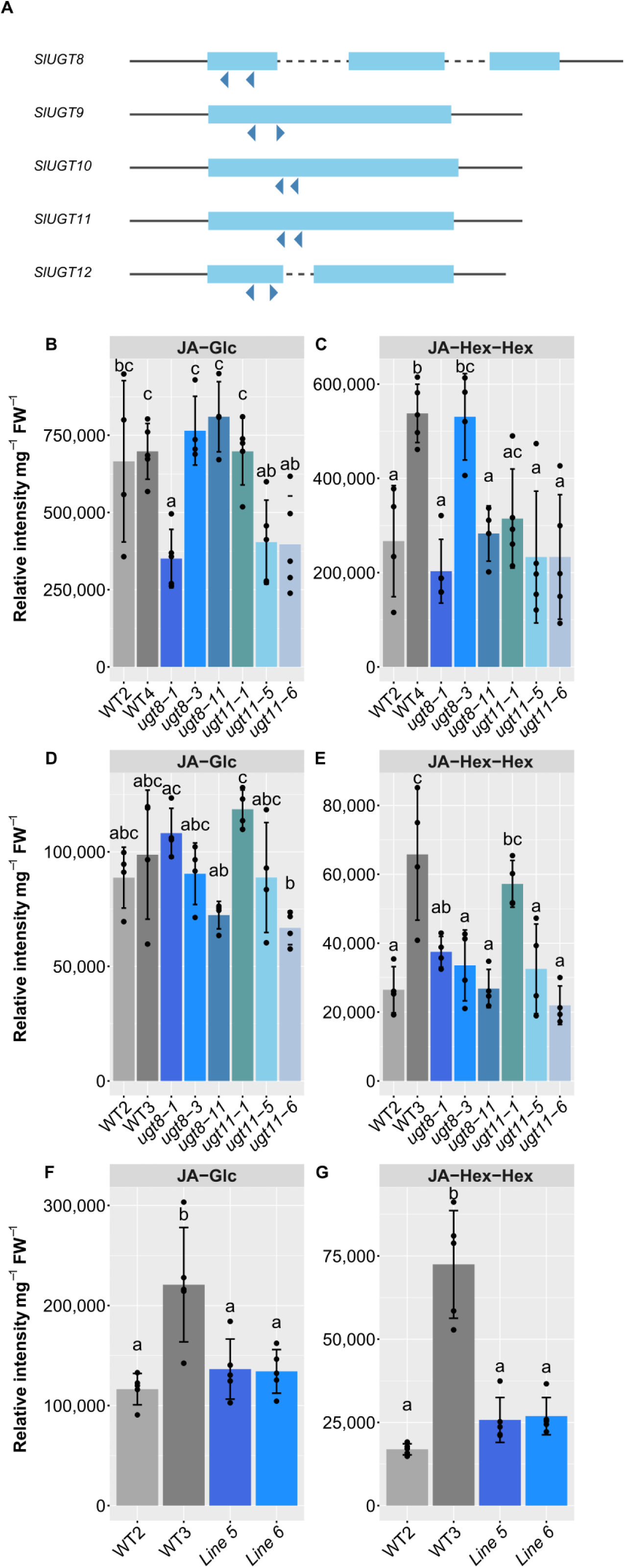
The analysis of metabolic phenotypes of tomato hairy root mutants. A. Schematics of five selected *SlUGT* genes that were targeted by two gRNAs to generate mutants. Light blue squares represent coding sequence exons, dashed lines introns, and blue arrow heads gRNAs. B-E. Two independent wild-type (WT), three *ugt8*, and three *ugt11* hairy root lines were treated with 50 µM JA for 6 h (B and C) or with 10 µM JA for 2 h (D and E). F-G. Two independent WT lines and two independent lines with gRNAs targeting all five *SlUGT*s (line 5 and line 6) were treated with 10 µM JA for 2h. Both JA-Glc (B, D, F) and JA-Glc-Glc (C, E, G) were measured. Black dots represent values for each biological replicate, bars represent the means, and error bars the standard deviation. Lower case letters represent statistical differences between treatments calculated by Tukey’s HSD test done following one-way ANOVA (p-value < 0.05).

Given that *SlUGT8* and *SlUGT11* encode enzymes capable of using JA as a substrate, we initially focused on single mutants of these genes (Supplementary Fig. 14) in our phenotypic analyses. If either of these genes is primarily responsible for JA glycosylation, we expect its loss-of-function mutant to exhibit a deficiency in JA glycosylation. To test this hypothesis, we treated tomato hairy root *SlUGT* mutants and WT lines with 50 µM JA for 6 h, as this treatment produced the highest levels of JA-Glc (Fig. 5A). Only one mutant line, *ugt8-1*, showed statistically significant difference from the WT lines in terms of JA-Glc levels (Fig. 8B). However, the two WT lines exhibited different levels of JA-Hex-Hex (Fig. 8C), suggesting that the lower levels of this metabolite in most mutant lines cannot conclusively be attributed to their genotypes. Since prolonged treatment with higher concentrations of JA may induce the expression of other *UGT*s capable of using JA as a substrate, potentially masking the phenotypes, we repeated the experiment with 10 µM JA treatment for 2 h (Fig. 8D and E). Under these conditions, no mutant was showed distinguishable differences from WT lines, suggesting that *ugt8* and *ugt11* single mutant lines are not deficient in JA glucosylation.

To explore whether *SlUGT8* and *SlUGT11* may be functionally redundant *in planta*, we generated lines in which all five selected *UGT*s were targeted. As SlUGT8 and SlUGT11 were shown to have JA glucosyltransferase activity (Fig. 4A), we aimed to isolate lines with mutations in at least these two genes. We identified two lines, designated as line 5 and line 6, both of which consistently carried mutations in both *SlUGT8* and *SlUGT11*. In line 5, both genes were 100% mutated, whereas in line 6, *SlUGT8* was fully mutated and *SlUGT11* was 53% mutated (Supplementary Table 4). *SlUGT9* was mutated in both lines (94% and 100% mutated in line 5 and line 6, respectively), while *SlUGT10* and *SlUGT12* were not mutated in line 6, and were 82% and 64% mutated in line 5, respectively. Due to the somatic nature of mutations in hairy roots, we were unable to isolate lines mutated in all five *UGT*s.

Having established a line with both *SlUGT8* and *SlUGT11* mutated, we treated lines 5 and 6 with 10 µM JA for 2 h. Both lines had comparable levels of JA-Glc (Fig. 8F) and JA-Hex-Hex (Fig. 8G) but were only statistically distinguishable from one independent WT line and not from the other. This indicates that the observed JA glucosylation levels are not solely attributable to their *ugt* genotypes. These findings further corroborate the hypothesis that SlUGT8 and SlUGT11 are unlikely to be the sole UGTs involved in JA-Glc biosynthesis in tomato.

## Discussion

### LiP-MS, a powerful tool for gene discovery in hormone homeostasis and signaling

In this study, using JA as the bait hormone and tomato as the model species, we demonstrated the effectiveness of LiP-MS as a tool to uncover new players in phytohormone homeostasis and signaling. LiP-MS is a relatively recent method for studying metabolite–protein interactions, and it is likely one of the most accessible and higher throughput options compared to other comparable proteomics techniques. In our LiP-MS experiments with varying JA concentrations, we detected known JA interactors, namely the tomato homolog(s) of Arabidopsis CYP20-3, a known JA-binding protein involved in light and redox signaling during stress responses^25^. Additionally, we identified several JA biosynthetic enzymes, including those catalyzing the final steps of the pathway, thus yielding JA as the direct product, as well as upstream enzymes. Altogether, these findings may point to the existence of hitherto unknown feedback mechanisms, which we did not further explore in this study. Finally, we discovered and validated two enzymes that glucosylate JA, likely constituting a previously unrecognized negative feedback loop in JA homeostasis and signaling.

### Glucosylation of JA as part of the JA endogenous responses

Our data indicate that JA glucosylation is an integral part of the JA response, at least in tomato, as we observed the accumulation of JA-Glc, the product of the UGTs identified by LiP-MS, following exogenous JA treatment and wounding-triggered endogenous accumulation of JA. The highest levels of JA-Glc were detected 6 h after exogenous JA treatment. Other reported JA-derivatives, such as 12-OH-JA and 12-HSO_4_-JA, have also been shown to be modulated by JA responses in tomato, and strikingly, both compounds exhibit peak accumulation at 6 h post-treatment^15^. Moreover, we found that the genes encoding the two validated UGTs were transcriptionally induced as early as 30 min after exogenous JA treatment, by which they form part of a transcriptional regulon, that includes not only genes known to be involved in the biosynthesis of JA, but also signaling components, such as several *JAZ* and *MYC* genes (Supplementary Fig. 15). These results align with the known feedback and amplification loops that modulate JA responses^5,6,9^.

### UGTs discovered by LiP-MS use JA as a substrate and catalyze the production of JA-Glc

We confirmed that at least two of UGTs identified by LiP-MS following JA treatment can catalyze reactions using JA as a substrate, producing JA-Glc. *In silico* analyses provided detailed metabolite–protein interaction 3D models and structural evidence for the interaction between JA and the selected UGTs. Notably, using AlphaFold and COACH-D modelling, we determined that the minimal atomic distances between atoms of JA and differentially abundant peptides from three of our lead UGTs (SlUGT8, SlUGT10, and SlUGT11) were within the 6.44 Å threshold, a distance suggested as a boundary of active sites^22^, indicating potential catalytic interactions. *In vitro* assays with recombinant versions of SlUGT8 and SlUGT11 confirmed their JA glucosylating activity, suggesting that this prediction method might be a reliable approach to prioritize candidates for functional validation. Notwithstanding, metabolite–protein interactions with minimal atomic distances greater than 6.44 Å could still indicate a different type of interactions, for instance, allosteric inhibition or regulation of protein complex stability. Therefore, this modeling information can guide the design of appropriate follow-up experimental characterization.

To our knowledge, no other UGTs that glycosylate JA to JA-Glc have been reported to date, making our findings an important step in elucidating a missing component of JA metabolism. Previous reports involving UGTs in JA responses have identified UGTs that glucosylate JA-derivatives. For instance, Haroth, et al. ^38^ identified Arabidopsis UGT76E1 and UGT76E2, both of which glucosylate 12-OH-JA. Likewise, the rice UGT, OsSGT, was found to glucosylate 12-OH-JA. Interestingly, OsSGT also glucosylates salicylic acid^39^. Notably, we discovered that SlUGT8 can catalyze the glucosylation of 12-OH-JA *in vitro*. Since many UGT enzymes are known to display varying degrees of substrate promiscuity^40,41^, it is possible that SlUGT8 and SlUGT11 may have affinities for other JA-derivatives as well. Because of the lack of available substrates, we were unable to explore this further.

The two other lead UGT proteins, SlUGT10 and SlUGT12, which we successfully produced recombinantly and whose corresponding genes were also JA-inducible, did not appear to be involved in the glucosylation of JA. However, it is possible that they use JA-Glc as a substrate to produce JA-Glc-Hex. This is possible, as we observed an increase in JA-Hex-Hex over time following exogenous application of JA, along with a concomitant decrease in JA-Glc.

Unfortunately, the tomato hairy root single and double mutants of *SlUGT8* and *SlUGT11* that we generated did not show reduced accumulation of JA-Glc or JA-Hex-Hex. It is plausible that only higher-order mutants would display a clear JA metabolic phenotype due to functional redundancy between these two and other, yet elusive, UGT enzymes. The enzymatic activities of SlUGT10 and SlUGT12 might not have been high enough to be detected under our experimental conditions, and since SlUGT9 could not be expressed in *E. coli*, we cannot exclude that these proteins glucosylate JA *in planta*. Moreover, there are still seven UGTs identified by LiP-MS that were not further investigated in this study, as they were not JA responsive at the transcriptional level. It cannot be excluded that some of these UGTs may also have JA-metabolizing functions, operating independently of the negative feedback loop, with constitutive basal expression levels sufficient to glucosylate (exogenous) JA. Generation of higher-order loss-of-function mutant plants is needed to test this hypothesis, as it would reduce the likelihood of obtaining chimeric and somatic mutations which often occur with multiplexed gene targeting in hairy roots.

### Glucosylation of JA as a potential negative feedback loop regulating JA homeostasis

We hypothesize that JA-Glc constitutes an inactive JA-derivative because one of the most well-established effects of JA, root growth inhibition^33–36^, was significantly reduced when we applied JA-Glc to tomato seedlings compared to JA. Accordingly, the effect on the expression of JA-inducible genes was also attenuated with JA-Glc treatment. Nevertheless, our experiments did not demonstrate that JA-Glc is completely inactive. JA-Glc treatment still impaired root growth and induced JA-responsive genes to some degree, albeit at an intermediate level between JA treatment (which resulted in pronounced root growth inhibition and high JA-inducible gene expression) and the control treatment (which showed no inhibition and low gene JA-inducible expression). One possibility is that the JA-Glc we synthesized in house and used in our assays was (partially) hydrolyzed in the medium, producing some free JA. Another possibility is that a proportion of the applied JA-Glc was converted back into JA within the plant by the action of specific, yet unknown, glucosidases. This was observed in tobacco cells treated with JA-Glc^32^, and a similar conversion has been observed for the glucosylated form of abscisic acid, another plant hormone^42^. Our observations are consistent with JA-Glc treaments of rice seedlings, where it had reduced effects on seedling growth compared to JA^43^. However, it is not known what the effects of JA-Glc are compared to no treatment, as such controls were not used by the authors. Independent of the results reported here, it is plausible to assume that JA-Glc is an inactive JA-derivative (or a less bioactive storage form of JA) because, to date, the only bioactive JA-derivatives identified involve conjugation of JA with hydrophobic amino acids such as JA-Ile, and the less bioactive JA-Leu, JA-Val, JA-Met and JA-Ala^11–13^. Other known JA-derivatives have not been classified as bioactive compounds and have not been found to bind to the JA receptor^2,44,45^. All of these bioactive conjugates are formed through the carboxyl group of JA, which is also the site of glucosylation in JA-Glc. This observation leads us to speculate that the formation of JA-Glc may deplete the pool of JA needed for JA-Ile biosynthesis.

Glucosylation of plant hormones is a common way to inactivate them. For example, the most abundant inactive auxin form is the indole-3-acetic acid-glucose conjugate (IAA-Glc). The glucosylation of auxins in Arabidopsis is performed by three UGTs, UGT84B1^46^, UGT74D1^47^, and UGT74E2^48^. Even though the preferred substrate of UGT74D1 is IAA, there is evidence indicating it can also use JA as a substrate to generate an uncharacterized product^49^. Since auxin and JA signaling pathways share high mechanistic similarities^50^, it is possible that they share metabolic regulators, potentially interconnecting auxin and JA signaling in a previously unknown way.

For brassinosteroids, the known UGTs rendering them inactive in Arabidopsis are UGT73C5^51^ and UGT73C6^52^. For cytokinins, two known Arabidopsis UGTs forming inactive glycoconjugates regulating homeostasis are UGT73C1 and UGT73C2^53,54^. For abscisic acid, UGT71C5 plays a major role in glucosylation, inactivation, and homeostasis^55^. For salicylic acid, three UGTs are known, UGT74F1, UGT74F2 and UGT76B1, the latter being the major active one^56,57^. These are only a select set of examples of Arabidopsis UGTs that illustrate that glucosylation is a crucial mechanism for controlling the physiological concentration of plant hormones. Glucosylation of plant hormones may modify their solubility and subsequently facilitate further processes like transport, storage, and degradation, ultimately maintaining hormonal homeostasis.

Upon compiling all evidence, our data suggest that the glucosylation of JA by SlUGT8 and SlUGT11 may be part of a fine-tuning mechanism that regulates JA hormonal homeostasis in response to stress. Eventually, JA glucosyl esters may be stored and later deglycosylated, suggesting that JA glucosylation may serve a dual function – regulating (active) JA pools and facilitating JA storage. As such, the UGTs involved in JA glycosylation may constitute a vital mechanism that regulates JA signaling, helping to maintain the stability and functioning of the JA system in an everchanging environment.

## Methods

### Generation of tomato hairy roots

Tomato hairy roots were generated by infecting cotyledons of tomato with *Agrobacterium (Rhizobium) rhizogenes,* as previously described in Ron, et al. ^58^ and Swinnen, et al. ^37^. Briefly, to generate control lines *Pro35S:GUS* in the Gateway vector pK7WG2D was introduced into *A. rhizogenes* strain ATCC15834 and used to infect cotyledons of tomato cv. MoneyMaker. Next, infected cotyledons were placed in nonselective Murashige and Skoog (MS) medium with 3% (w/v) sucrose. After three days, the cotyledons were moved to MS medium containing 200 µg/mL of cefotaxime and 50 µg/mL of kanamycin. Following two weeks of incubation in a dark environment at 22-25°C, new roots developed from the cotyledons. These newly formed roots were detached and transferred to selective medium. The process of transferring roots was repeated twice at three weeks interval. After six weeks of growth on selective medium, the roots were moved to a nonselective medium and used in LiP-MS experiments. When the root material was needed for more experiments, tomato hairy root cultures were sub-cultured onto new plates containing nonselective media every 4-5 weeks.

### LiP-MS experiments

The LiP-MS experiments were performed according to the published detailed protocol^10^, also described below.

#### LiP-MS sample preparation

LiP-MS experiments were conducted on samples of tomato hairy root cultures using four biological replicates for each construct. Briefly, samples were ground in liquid nitrogen and then suspended in extraction buffer (100 mM HEPES pH 7.5, 150 mM KCl, 1 mM MgCl_2_) at a 1:1 weight-to-volume ratio (w/v). The homogenized suspensions were subjected to three rounds of mechanical freeze-thaw lysis to achieve complete cell lysis. The resulting lysates were sonicated using a Bioruptor™ Next Gen sonicator at level 5 for twenty cycles of 10 s on, and 10 s off. Next, the samples were centrifuged at 20,000 × g for 10 min at 4°C to remove cell debris, and the resulting supernatants were carefully transferred to new tubes. These centrifugation and transfer steps were repeated once. Then, protein lysates were desalted using NAP™-5 20 ST columns following the manufacturer’s instructions. Next, the protein concentration for each sample was determined using the Bradford assay. Then, the lysates from independent biological replicates were aliquoted into equivalent volumes containing 100 µg of total protein in 1.5-mL Eppendorf tubes. Then, samples were pre-incubated at 25°C for 5 min in a dry block incubator. Next, the metabolite solution or the corresponding solvent for mock was added to the samples and was then incubated for 10 min at 25°C. Subsequently, proteinase K was added to the samples at an enzyme-to-substrate (E:S) ratio of 1:100 (w/w), and the mixture was incubated for 5 min at 25°C. The samples were heated at 98°C for 3 min to denature proteinase K and stop proteolytic digestion. Next, overnight digestion was performed using mass spec grade trypsin/Lys-C Mix at an E:S ratio of 1:50 (w/w) at 37°C. The next day, the samples were acidified by adding trifluoroacetic acid (TFA) 5% to reach a final concentration of 0.5% in each sample. The samples were then centrifuged at 20,000 × g and 4°C for 10 min to remove insoluble particulates, and the resulting supernatants were transferred to new tubes. The centrifugation and transfer steps were repeated once. Next, methionine oxidation was performed by adding 30% (v/v) hydrogen peroxide to each sample to a final concentration of 0.5%, and the samples were incubated for 30 min at 30°C while mixing. C18 reversed-phase sorbent pipette tips (Bond Elut OMIX 100 µL C18 tips) were used according to the manufacturer’s instructions for solid-phase extraction of peptides. The samples were then vacuum-dried using a SpeedVac concentrator for 4 h or overnight at 30-40°C, followed by reconstitution in 30 µL of 2-mM TCEP in water: acetonitrile 98:2 v/v. Finally, the samples were subjected to shotgun mass spectrometry data acquisition.

#### LiP-MS mass spectrometry data acquisition

We performed LC-MS/MS shotgun analysis for a long 3-h LC-run using a Q-exactive-HF mass spectrometer, Thermo Fisher Scientific (for DDA proteomics), using in-house settings. Specific details about the MS settings are described in Venegas-Molina, et al. ^10^.

#### LiP-MS data analysis

Briefly, DDA raw proteomics data from LiP-MS experiments was mapped against the UniProt protein database of tomato using the MaxQuant software, applying a semi-tryptic digestion rule. Resulting LFQ intensities were used for filtering, data processing and statistical analysis using the software Perseus^59^. The filtering step included filtering contaminants and decoys. For data analysis only unique peptides were selected, following log transformation, normalization, and data imputation as integrated in Perseus. Lastly, a differential enrichment statistical analysis was performed. For this, at least three biological replicates in at least one treatment were statistically tested to detect differential peptide abundance between treatments using the *t*-test with Benjamini-Hochberg FDR correction integrated into Perseus. Volcano plots and Venn diagrams were drawn with the resulting data. Retention was calculated as the percentage of protein interactors detected from the lowest concentration retained at the highest concentration.

### Liquid chromatography-mass spectrometry (LC-MS) targeted metabolomics experiment for measuring JA-Glc and JA-Hex-Hex

About 1 cm fresh root tips of tomato hairy root cultures grown on solid medium were inoculated into 5 mL liquid MS medium supplemented with vitamins and 3% (w/v) sucrose for two weeks in 50-mL conical tubes (Falcon) with shaking at 130 rpm on platform shakers. After one week, additional 5 mL of medium was added to each culture. Samples were treated by replacing medium with 10 mL fresh medium with JA (diluted in ethanol), or mock (equal volume of ethanol). Samples were harvested after 3, 6, 12, or 24 h and stored at −70°C until metabolite extraction. For leaf wounding experiments, third leaves of 5-week-old Money Maker plants (biological replicates) were wounded along the entire surface using wide forceps. Samples were harvested after 0, 1, and 3 h and stored at −70°C until metabolite extraction. The metabolite extraction from plant material consisted of grinding samples on a tissue homogenizer (Retsch) for 1 min at 20 Hz in 2-mL microcentrifuge tubes with two 4-mm steel beads. Next, the ground powder was transferred to new 2-mL microcentrifuge tubes, weighted and fresh weight recorded. Next, 1000 µL of MeOH (HPLC grade) was added to each tube. Samples were shaken for 1 h at 4°C, then centrifuged at 16 000 *g* for 10 min. 800 µL of methanolic extracts was transferred to 2-mL microcentrifuge tubes. For removal of lipids, C18 SPE cartridges (Waters catalog number WAT094225) were used following the manufacturer’s instructions. The SPE columns were conditioned with 0.5 mL MeOH, the samples were added, collected into a 2-mL tube, and the cartridge was washed twice with 400 µL MeOH into the same tube. Next, the samples were evaporated in a vacuum centrifuge (LabConco). Then, the samples were resuspended in 100 µL of LC-MS-grade water, transferred to 96-well 0.2-µm filter plates on top of 96-well plates, and collected after centrifugation.

Samples were subjected to Ultra Performance Liquid Chromatography High Resolution Mass Spectrometry (UPLC-HRMS) at the VIB Metabolomics Core Ghent (VIB-MCG). 10 µl was injected on a Waters Acquity UHPLC device connected to a Vion HDMS Q-TOF mass spectrometer Chromatographic separation was carried out on an ACQUITY UPLC BEH C18 (50 × 2.1 mm, 1.7 μm) column from Waters, and temperature was maintained at 40°C. A gradient of two buffers was used for separation: buffer A (99:1:0.1 water:acetonitrile:formic acid, pH 3) and buffer B (99:1:0.1 acetonitrile:water:formic acid, pH 3), as follows: 99% A for 0.1 min decreased to 50% A in 10 min. The flow rate was set to 0.5 mL min^−1^. Negative Electrospray Ionization (ESI) was applied in TOF-MRM mode. The LockSpray ion source was operated in negative electrospray ionization mode under the following specific conditions: capillary voltage, 2.5 kV; reference capillary voltage, 2.5 kV; source temperature, 120°C; desolvation gas temperature, 600°C; desolvation gas flow, 800 L h^−1^; and cone gas flow, 50 L h−1. The collision energy was set at 30eV for each transition. Nitrogen (greater than 99.5%) was employed as desolvation and cone gas. Leucine-enkephalin (250 pg μL^−1^ solubilized in water:acetonitrile 1:1 [v/v], with 0.1% formic acid) was used for the lock mass calibration, with scanning every 2 min at a scan time of 0.1 s. Profile data was recorded through Unifi Workstation v2.0 (Waters).

For MS, ion counts of [M-H+FA] were measured for JA-Glc and JA-Hex-Hex. A chemically synthesized JA-Glc was used as an external standard.

### Chemical synthesis of JA-Glc

The chemical synthesis of JA-Glc has been reported by Qian et al.^60,61^, in a three-step approach starting from methyl jasmonate (MeJA). There, MeJA was hydrolyzed to the free carboxylic acid, then activated as the acid chloride and finally coupled to D-glucose in an acylation reaction. As MeJA is a mixture of isomers, and the acylation can proceed on multiple hydroxyls, this approach leads to a rather complex mixture of JA-Glc isomers. For our synthesis of JA-Glc, we started from racemic (+/-) JA (Duchefa: J0936.0250) and adopted a novel chemo- and regioselective carboxylic acid glycosylation protocol reported by Takeuchi et al.^61^. The one-step Mitsunobu-type coupling of JA and α-D-glucose afforded JA-Glc in a much purer form, consisting of only two main stereoisomers (∼90%), both showing the inverted β configuration. The remaining main 50:50 stereoisomerism is directly related to the racemic JA starting material (which is also used for the control treatments with JA). This novel approach also allowed the characterization of the synthetic JA-Glc via 1D and 2D NMR spectroscopy^60,62,63^. The full NMR data, appended with supporting information for the synthesis of JA-Glc have been included in Supplementary dataset 3.

### Root growth inhibition bioassay

Tomato seeds cv. Moneymaker were germinated and grown on solid MS medium with 1% (w/v) sucrose and 1% (w/v) agar for one week, and immediately transferred to plates containing the same medium supplemented with either 1 µM JA, 1 µM JA-Glc, or mock (equivalent volume of ethanol). After two or three days of growth on plates with treatments, roots of the seedlings were imaged with a scanner and measured using Fiji^64^. Fifteen plants (biological replicates) were used per experiment, grown on three separate plates. Statistical analysis and plotting were done using R through RStudio. Results of statistical tests are available in Supplementary Dataset 4.

### 3D modeling and structure analysis of metabolite–protein interactions

We used AlphaFold to obtain 3D structures of the proteins of interest^29^ by either retrieving them from the AlphaFold Protein Structure Database (AlphaFold DB, https://alphafold.ebi.ac.uk) when available^65^ or predicting them using AlphaFold installed in the High-Performance Computing infrastructure of Ghent University (HPC-UGent) (Tier-2 system via SSH protocol). Next, we obtained the 3D structures of the metabolites of interest from the PubChem database^30^. Then, we applied molecular docking through the COACH-D platform^31^ to construct 3D structure models of the metabolite–protein interactions. The COACH-D algorithm first predicts binding pockets of an input protein and then uses the three-dimensional structure of a metabolite of interest to predict the binding poses with higher binding affinity and the preferred orientation of the metabolite into the protein binding pockets using molecular docking. Finally, it builds a resulting complex 3D structure. For each prediction, COACH-D generates a confidence score (C-score) according to binding affinity to judge the reliability of each prediction. The C-score ranges from 0 to 1, with higher scores indicating more reliable predictions. Next, using the 3D structure models, we compared the positions of the peptides detected with LiP-MS versus the JA-UGT binding sites predicted with COACH-D. To obtain a higher sequence coverage, in the case of the five selected UGTs, we retrieved all the peptides identified in LiP-MS with 10 mM JA, selected the ones that passed the statistical threshold cut-off q-value < 0.05, and mapped them in the 3D metabolite–protein structures. Finally, we measured the atomic distances between atoms of the metabolite of interest and differentially abundant peptides, calculated minimal atomic distances in angstroms (Å), and compared them with distances of known catalytic interactions using the software Yasara (yasara.org).

### Expression of UGT proteins

The coding sequence of each UGT of interest was cloned using the Gateway cloning method after amplification with primers listed in Supplementary Dataset 5, first in the donor vector pDONR221 and subsequently in the destination vector pET-DEST42 to produce SlUGT proteins fused at their C-terminus to a 6xHis-tag. The expression clones were transformed into *E. coli* Rosetta (DE3) strain and tested for protein expression and solubility. For expression tests, *E. coli* precultures were grown in LB media supplemented with antibiotics (chloramphenicol and carbenicillin) at 37°C, 200 rpm overnight. The next day, three separate cultures were prepared by inoculating 150 μL of the overnight preculture in 25 mL of LB supplemented with antibiotics and incubated at 37°C, 200 rpm until reaching an optical density (OD600) of ∼0.6. Then, the cultures were induced with 1 mM IPTG and incubated at 20°C, 28°C, or 37°C, at 200 rpm. After 3 and 20 h, an aliquot of 4 μL for each condition was collected and assessed by sodium dodecyl-sulfate polyacrylamide gel electrophoresis (SDS-PAGE) and Western blotting (WB) using a Penta-His HRP conjugate antibody (Qiagen). The growth conditions that showed a visible protein with the expected molecular weight were used for solubility tests. For solubility tests, pellets selected from the expression tests were weighed and resuspended in buffer 1 (300 mM NaCl, 50 mM Tris, 20 mM imidazole, and protease inhibitors without EDTA, pH to 7.5) and sonicated for 2 min (10 s on, 20 s off, 40% amplitude) with a probe sonicator. Then, samples were centrifugated for 10 min (14 000 *g*) at 4°C. The supernatants were collected, and the pellets were stored; additionally, a 60-μL subsample of each supernatant was taken. Next, His Mag Sepharose Ni magnetic beads (Sigma-Aldrich) were equilibrated using buffer 2 (300 mM NaCl, 50 mM Tris, and protease inhibitors EDTA Free, pH to 7.5), placed in a magnetic stand, and the supernatant removed. Then, samples were resuspended with beads and incubated on a rotating wheel for 25 min. The beads were separated, and the supernatant was removed. The beads were washed four times with buffer 1. A subsample of 60 μL of the first washing step was collected. Next, beads were eluted three times with buffer 3 (300 mM NaCl, 50 mM Tris, 500 mM imidazole, and protease inhibitors without EDTA, pH to 7.5), and each fraction was collected. The different eluted fractions together with aliquots of the input, pellet’s debris, and washing steps were used for SDS-PAGE and WB using anti-His-HRP antibody. The conditions showing clear bands of the proteins with the expected molecular weights from the elution fractions were selected for big-scale protein purification.

### Purification of SlUGT proteins

Proteins were purified by a two-step protein purification strategy of immobilized metal affinity chromatography (IMAC) and size exclusion chromatography (SEC). Briefly, the best conditions from the expression and solubility tests were selected for large-scale expression and purification of SlUGT proteins. *E. coli* cultures were prepared using the selected conditions for each SlUGT protein but in a volume of 6 L. Next, the cultures were centrifuged, and the pellets were collected, weighed, and stored at −70°C. Pellets were resuspended in extraction buffer (300 mM NaCl, 50 mM Tris/HCl pH 7.5, 20 mM imidazole, 1 mM PMSF, pH to 7.5, ratio 3 mL buffer/mg pellet), sonicated for 10 min (10 s on; 10 s off; 40% amplitude) and centrifuged for 30 min (20 000 *g* at 4°C). The supernatants were filtered and immediately loaded onto the ÄKTA chromatography system for IMAC. After IMAC, resulting elution fractions together with aliquots of input, flow-through, and washing steps were analyzed by SDS-PAGE and WB using anti-His HRP antibody. Five elution fractions containing the protein of interest were selected according to the results of SDS-PAGE, WB, and chromatograms; pooled, and delivered for a SEC run to obtain cleaner proteins. During SEC, proteins were eluted in a buffer containing 150 mM NaCl, 50 mM Tris/HCl pH7.5, pH to 7.5. After SEC, resulting elution fractions were subjected to SDS-PAGE and WB with anti-His HRP antibody. Elution fractions showing the protein of interest at the expected molecular size were selected according to results of SDS-PAGE, WB, and chromatograms, pooled and concentrated using 30 kDa MWCO Centricon centrifugal filter units (Millipore). Finally, protein concentrations were measured using the Bradford method, and aliquots of each SlUGT protein were stored at −70°C.

### LC-MS-based activity assays

Activity assays were performed by incubating 0.5 mM JA and 0.5 mM UDP-glucose as substrates with 20 µg of each purified SlUGT-6xHis (SlUGT8, SlUGT10, SlUGT11, SlUGT12) to a final volume of 100 µL and incubated for 30 min or overnight at room temperature. The reactions were stopped by adding 50 µL MeOH. As a control, we used the same experimental setup, but the SlUGT proteins were boiled before the assay to inactivate them by denaturation. The resulting reaction was injected and monitored by high-resolution UHPLC-QTOF-MS^66^: UHPLC1290 Infinity (Agilent Technologies, Santa Clara, CA, USA) coupled to an HRMS instrument (6540 or 6546 UHD Accurate-Mass Q-TOF, Agilent Technologies, Santa Clara, CA, USA) with Agilent Dual Jet Stream Technology as electrospray ionization (ESI) source (Agilent Technologies, Santa Clara, CA, USA).

### Glycosyltransferase activity assays

We used the commercial kit UDP-Glo™ Glycosyltransferase Assay (Promega; catalog number V6961) and followed the manufacturer’s instructions. To determine the *in vitro* minimal concentration of SlUGT8 protein for JA glucosylation, we incubated 50 µM JA and 100 µM UDP-glucose with 111, 333, and 1000 ng of purified SlUGT8-6xHis. As control, we used the same experimental setup, but the SlUGT8 protein was boiled before the assay. In this assay, we also included controls samples with only SlUGT8, and with only SlUGT8 and JA. For testing the sugar donor specificity of SlUGT8, we incubated 50 µM JA with either 100 µM UDP-glucose or UDP-glucuronic acid, together with six different concentrations of SlUGT8 protein. As control, we used the same experimental setup, but SlUGT8 was boiled before the assay. In this assay, we also included controls samples with only SlUGT8, and with only SlUGT8 and JA.

### Gene expression analyses

Total RNA from tomato roots (cv. Moneymaker) was extracted and purified with the ReliaPrep® RNA Tissue Miniprep System (Promega) according to manufacturer’s instructions. The Superscript cDNA Synthesis Kit (Quantabio) was used for cDNA synthesis from 1 µg of RNA. qRT-PCR was performed using diluted cDNA (1:8) on a LightCycler 480 (Roche Diagnostics) in 384-well plates with LightCycler 480 SYBR Green I Master (Roche) according to the manufacturer’s instructions. Expression levels of target genes were normalized relative to Solyc10g049850 (SlTIP41). Primers used for RT-qPCR are listed in Supplementary Dataset 5. Analysis of relative gene expression data was performed using the 2−^ΔΔ^CT method. The data statistics were calculated through Tukey’s HSD test, following one-way ANOVA. Because of the dynamic range of JA-induced gene expression, the obtained values did not always follow a normal distribution, as indicated by the D’Agostino-Pearson test. The non-normally distributed expression values were log2-transformed to obtain a normal distribution before ANOVA analysis.

### Generation of *SlUGT* tomato hairy root mutant lines

We generated UGT loss-of-function lines following the protocol by Swinnen, et al. ^37^. Briefly, coding sequences of *SlUGT8, SlUGT9, SlUGT10, SlUGT11*, and *SlUGT12* were retrieved from the SolGenomics Network database^67^. Next, two guide RNAs were designed for each *SlUGT* (Supplementary Dataset 5) targeting sequence regions corresponding to the first exon of each gene with the aim to generate a big deletion visible on gel to facilitate genotyping. We used the Golden Gate method to generate CRISPR constructs that consisted of the *Petroselinum crispum* UBIQUITIN (*PcUBI*) promoter, *AtCas9* with A2 sequence and *mCherry*, and two *AtU6*-gRNAs specific for each *SlUGT* in a pFASTRK backbone. We used the vector ID: 12_11 from the VIB Gateway collection (pFASTRK-PcUBIP-AtCas9-NLS-P2A-mCherry-G7T-A-CmR-ccdB-G). Constructs were transformed into *A. rhizogenes* strain ATCC15834, and cotyledons of tomato seedlings were transformed as described above. We generated 40 different tomato CRISPR hairy root lines per *SlUGT*. Lines were subcultured every two weeks. After three runs of culturing, roots were genotyped to verify the mutations generated. Plant issue was collected on QIAcard™ FTA™ PlantSaver cards (Qiagen) by pressing through Parafilm®. Genotyping was done by Inference of CRISPR Edits (ICE) analysis (Synthego; ice.synthego.com) using Sanger-sequenced purified amplicons amplified using primers listed in Supplementary Dataset 5. PCR amplifications were done on 1.2-mm FTA card punches washed according to manufacturer’s instructions. Amplicons were purified using HighPrep™ PCR cleanup system (MagBio), and Sanger sequenced using amplification primers. For each *SlUGT*, we selected the best four putative loss-of-function tomato lines according to their mutated genotype and homozygosity after at least two rounds of genotyping.

## Data, Materials, and Software Availability

All data are included in the article, SI and/or public databanks. RNA-Seq data have been deposited in the ArrayExpress database (accession E-MTAB-11876 and E-MTAB-14610). The MS data from the LiP-MS experiments have been deposited via ProteomeXchange in the PRIDE database (accession PXD057493).

## Acknowledgements

We thank Annick Bleys for critically reading and help with preparing the manuscript, Nikolaos Ntelkis for help in submitting the RNA-Seq data, the VIB Proteomics Core Facility for running proteomics samples on LC-MS, and the VIB Metabolomics Core Facility (Ghent) for running metabolite analyses. This research was supported by the European Union’s Horizon 2020 research and innovation program under Grant Agreements No 825730 (Endoscape) and No 101000373 (InnCoCells) and the Research Foundation Flanders (FWO) through the projects G008417N and G004320N, all assigned to A.G.. P.V.D. was supported by the H2020 European Research Council (ERC) under the European Union’s Horizon 2020 research and innovation program (PROPHECY grant agreement No 803972). K.Š., S.S.G. and A.C.J.-M. were supported by postdoctoral fellowship from the European Union’s Horizon 2020 research and innovation program under the Marie Skłodowska-Curie grant agreement No 101151286, the Research Foundation Flanders (FWO), and the Special Research Fund from Ghent University, respectively. We thank the UGent NMR core facility for 700 MHz spectroscopy, which is funded by the Hercules foundation (AUGE/15/2012) and FWO (I009324N). J.M.W and H.J. thank BOF UGent for funding. L.M. was supported by the Goettingen Graduate School for Neurosciences, Biophysics, and Molecular Biosciences in frame of the PRoTECT program at the Georg August University Goettingen. I.F. acknowledges funding from the Deutsche Forschungsgemeinschaft (GRK 2172-PRoTECT, INST 186/1434-1, and ZUK 45/2010).

## Author contributions

A.G. initiated and managed the project, designed experiments, analyzed data, and wrote the manuscript; J.V.-M., and K.Š. designed and performed experiments, analyzed data, and wrote the manuscript; L.M., S.S.G., H.J., A.C.J.-M., E.L., and P.V.D. designed and performed experiments and analyzed data; J.W. and I.F. designed experiments and analyzed data; all authors discussed the results and commented on the manuscript.

