## Supplemental Figures and Tables for "Discovery of tomato UDP-glucosyltransferases involved in bioactive jasmonate homeostasis using limited proteolysis-coupled mass spectrometry"

### Supplemental Information

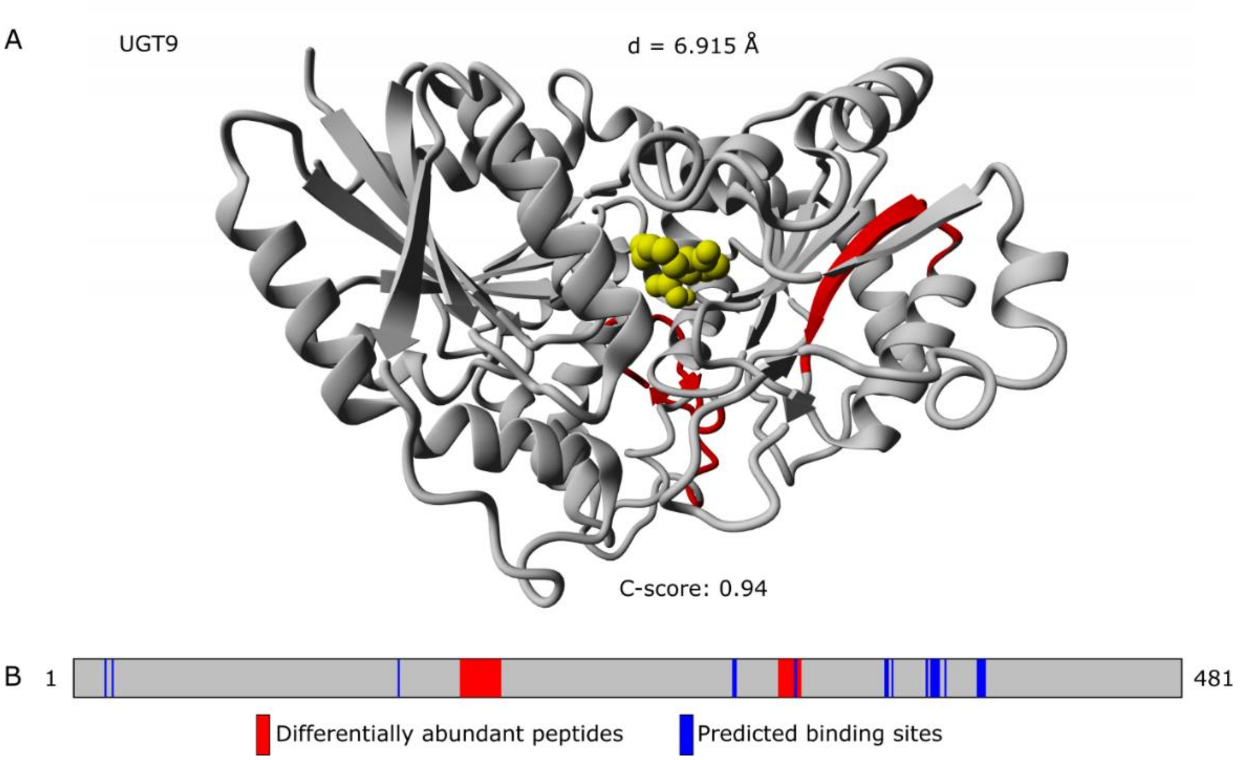


**Supplementary Fig. 1. 3D metabolite–protein interaction structural model of JA–SlUGT9**. A. 3D structural model showing the interaction between JA (yellow) and SlUGT9 (gray), with differentially abundant peptides detected by LiP-MS highlighted in red. B. Protein map illustrating the overlap between the differentially abundant peptides detected by LiP-MS (red) and the COACH-D predicted binding sites (blue). C-score: COACH-D model confidence score as defined by Wu, et al. [1]. d: minimal distance between the atoms of JA and the differentially abundant peptides detected by LiP-MS

A

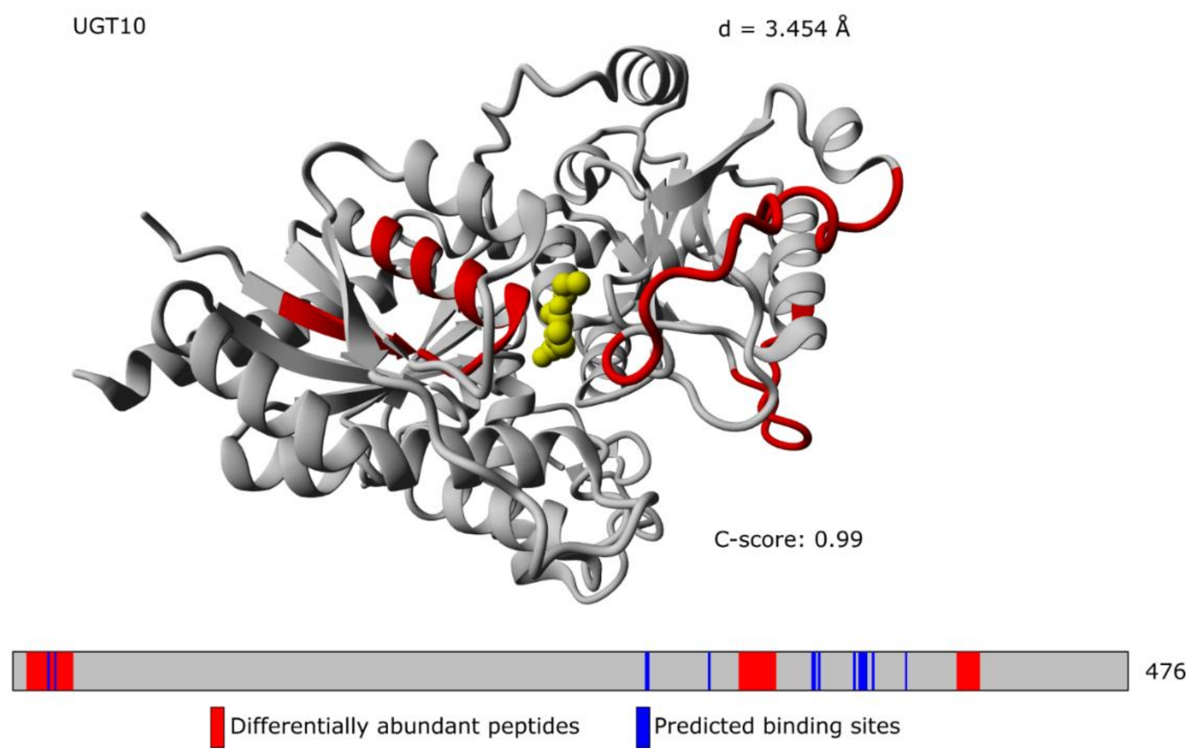


**Supplementary Fig. 2. 3D metabolite–protein interaction structural model of JA–SlUGT10**. A. 3D structural model showing the interaction between JA (yellow) and SlUGT10 (gray), with differentially abundant peptides detected by LiP-MS highlighted in red. B. Protein map illustrating the overlap between the differentially abundant peptides detected by LiP-MS (red) and the COACH-D predicted binding sites (blue). C-score: COACH-D model confidence score as defined by Wu, et al. [1]. d: minimal distance between the atoms of JA and the differentially abundant peptides detected by LiP-MS.

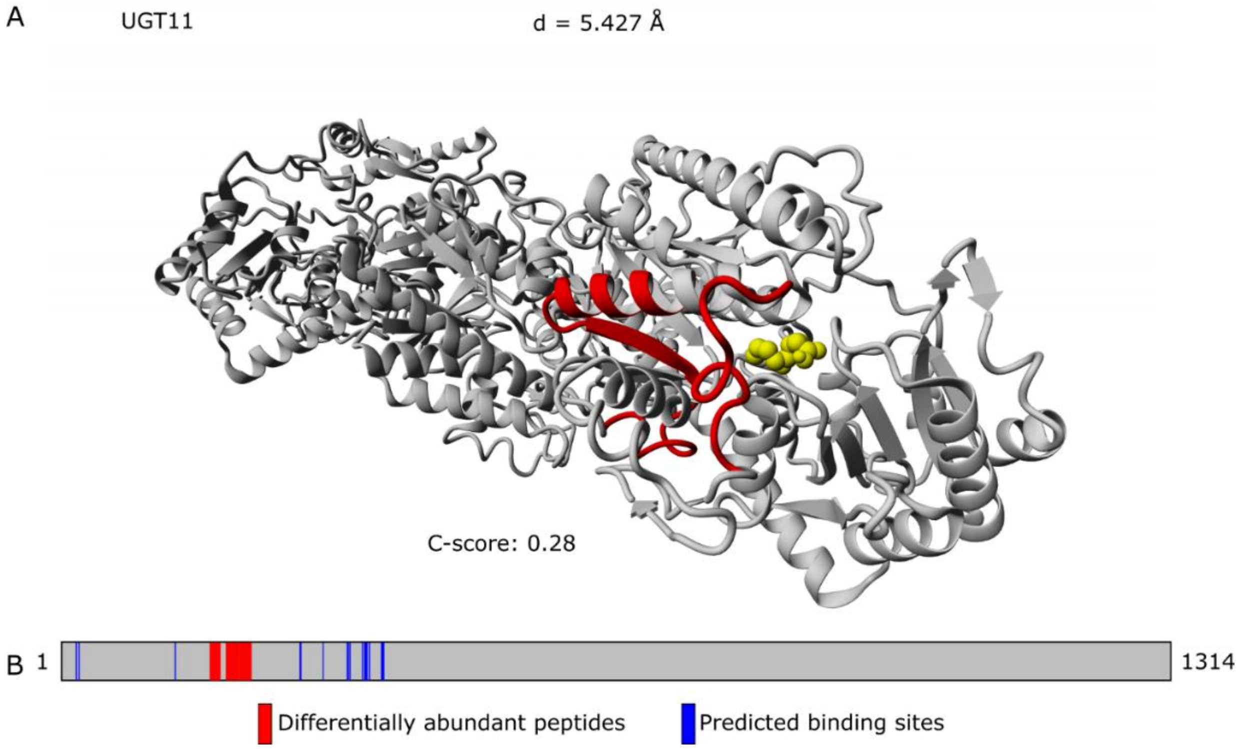


**Supplementary Fig. 3. 3D metabolite–protein interaction structural model of JA–SlUGT11**. A. 3D structural model showing the interaction between JA (yellow) and SlUGT11 (gray), with differentially abundant peptides detected by LiP-MS highlighted in red. B. Protein map illustrating the overlap between the differentially abundant peptides detected by LiP-MS (red) and the COACH-D predicted binding sites (blue). C-score: COACH-D model confidence score as defined by Wu, et al. [1]. d: minimal distance between the atoms of JA and the differentially abundant peptides detected by LiP-MS.

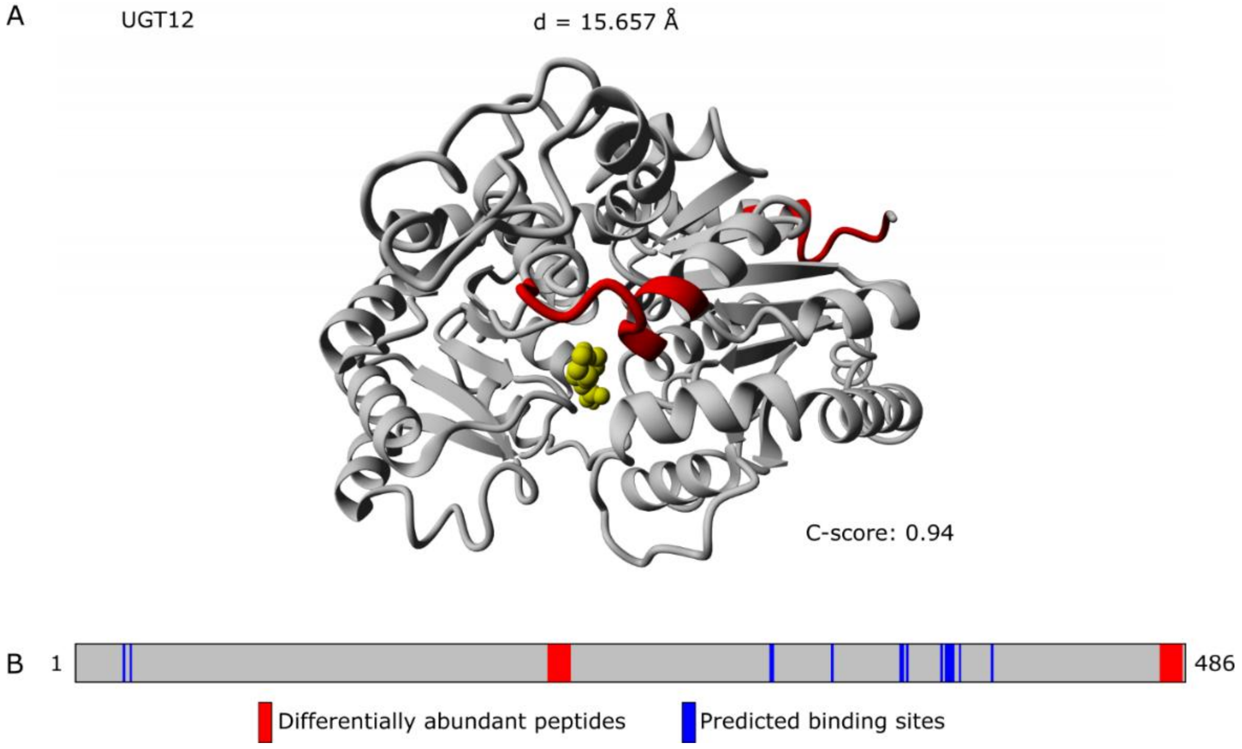


**Supplementary Fig. 4. 3D metabolite–protein interaction structural model of JA–SlUGT12**. A. 3D structural model showing the interaction between JA (yellow) and SlUGT12 (gray), with differentially abundant peptides detected by LiP-MS highlighted in red. B. Protein map illustrating the overlap between the differentially abundant peptides detected by LiP-MS (red) and the COACH-D predicted binding sites (blue). C-score: COACH-D model confidence score as defined by Wu, et al. [1]. d: minimal distance between the atoms of JA and the differentially abundant peptides detected by LiP-MS.

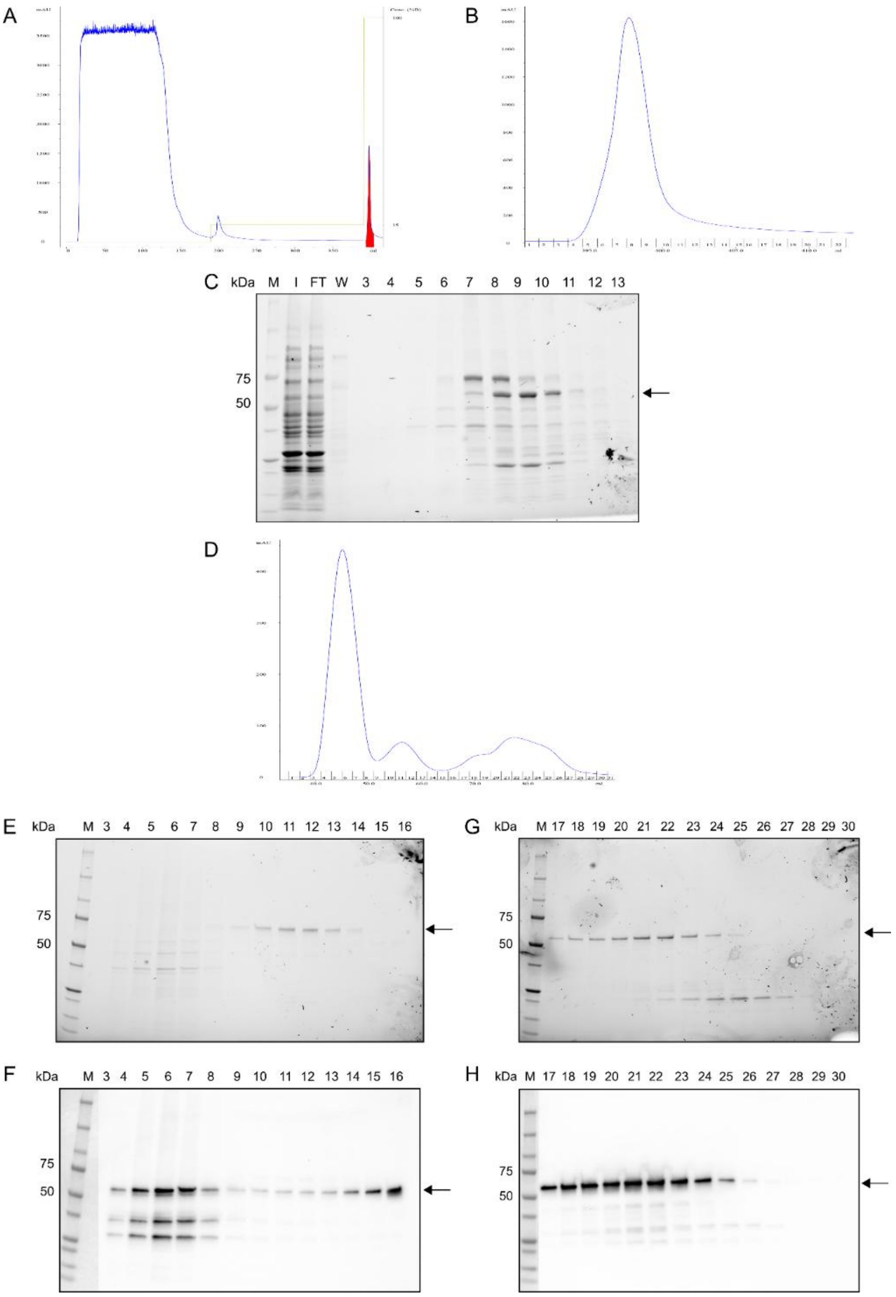


**Supplementary Fig. 5. Protein purification of SlUGT8.** SlUGT8 was fused to a His-tag, heterologously expressed in *E. coli* Rosetta (DE3), and purified by a two-step protein purification strategy of immobilized metal affinity chromatography (IMAC) and size exclusion chromatography (SEC). A. IMAC chromatogram. The chromatogram illustrates the absorption at 280 nm during elution (milli absorption units, mAU). The second y-axis shows elution buffer concentration (%). The fraction that contains the protein of interest is marked in red. B. IMAC chromatogram showing a zoom-in view of the peak containing the protein of interest. The x-axes show volume (mL) and IMAC fractions. The IMAC fractions were later used for sodium dodecyl sulfate (SDS)-polyacrylamide gel electrophoresis (PAGE). C. SDS-PAGE of IMAC fractions with samples of input (I), flow-through (FT), wash (W), and fractions 3 to 13. The black arrow highlights the size of the recombinant UGT8 protein at 58.1 kDa. Later, selected IMAC fractions were combined for SEC. D. SEC chromatogram. The x-axes show volume (mL) and SEC fractions. E. SDS-PAGE of SEC fractions 3 to 16. F. Western blot (WB) of SEC with anti-His-HRP antibody fractions 3 to 16. G. SDS-PAGE of SEC fractions 17 to 30. H. WB of SEC with anti-His-HRP antibody fractions 17 to 30.

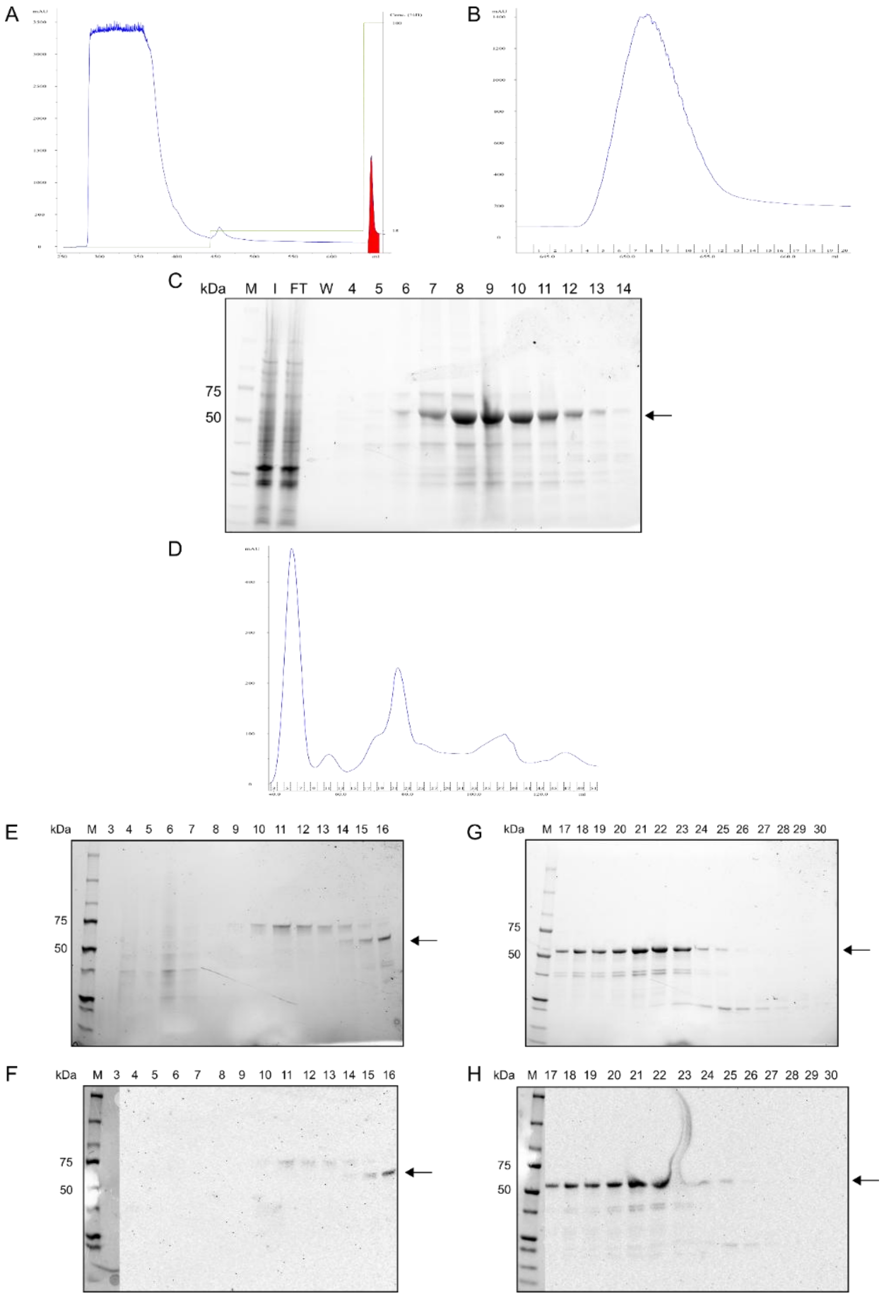


**Supplementary Fig. 6. Protein purification of SlUGT10.** SlUGT10 was fused to a His-tag, heterologously expressed in *E. coli* Rosetta (DE3), and purified by a two-step protein purification strategy of immobilized metal affinity chromatography (IMAC) and size exclusion chromatography (SEC). A. IMAC chromatogram. The chromatogram illustrates the absorption at 280 nm during elution (milli absorption units, mAU). The second y-axis shows elution buffer concentration (%). The fraction that contains the protein of interest is marked in red. B. IMAC chromatogram showing a zoom-in view of the peak containing the protein of interest. The x-axes show volume (mL) and IMAC fractions. The IMAC fractions were later used for sodium dodecyl sulfate (SDS)-polyacrylamide gel electrophoresis (PAGE). C. SDS-PAGE of IMAC fractions with samples of input (I), flow-through (FT), wash (W), and fractions 4 to 14. The black arrow highlights the size of the recombinant UGT10 protein at 56.8 kDa. Later, selected IMAC fractions were combined for SEC. D. SEC chromatogram. The x-axes show volume (mL) and SEC fractions. E. SDS-PAGE of SEC fractions 3 to 16. F. Western blot (WB) of SEC with anti-His-HRP antibody fractions 3 to 16. G. SDS-PAGE of SEC fractions 17 to 30. H. WB of SEC with anti-His-HRP antibody fractions 17 to 30.

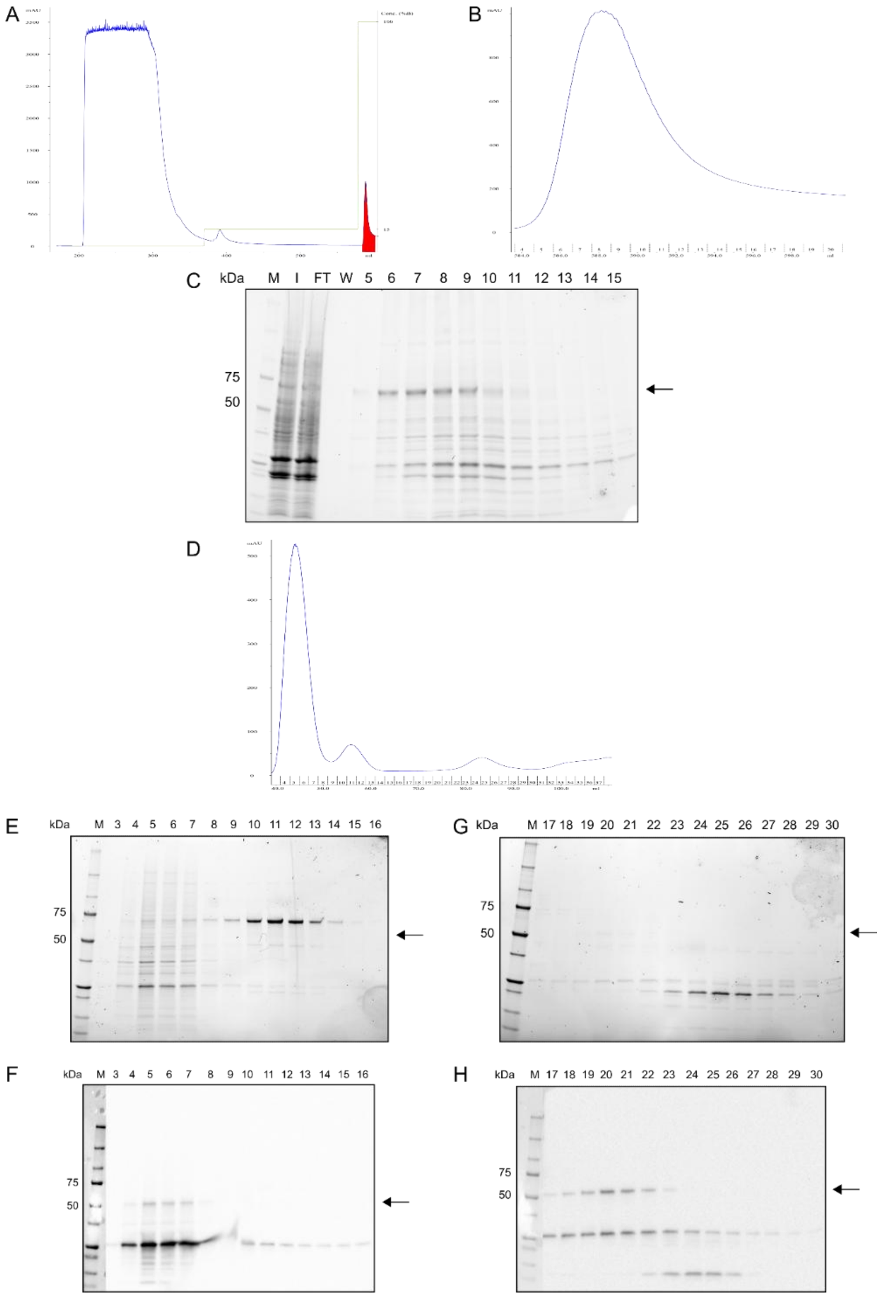


**Supplementary Fig. 7. Protein purification of SlUGT11.** SlUGT11 was fused to a His-tag, heterologously expressed in *E. coli* Rosetta (DE3), and purified by a two-step protein purification strategy of immobilized metal affinity chromatography (IMAC) and size exclusion chromatography (SEC). A. IMAC chromatogram. The chromatogram illustrates the absorption at 280 nm during elution (milli absorption units, mAU). The second y-axis shows elution buffer concentration (%). The fraction that contains the protein of interest is marked in red. B. IMAC chromatogram showing a zoom-in view of the peak containing the protein of interest. The x-axes show volume (mL) and IMAC fractions. The IMAC fractions were later used for sodium dodecyl sulfate (SDS)-polyacrylamide gel electrophoresis (PAGE). C. SDS-PAGE of IMAC fractions with samples of input (I), flow-through (FT), wash (W), and fractions 5 to 15. The black arrow highlights the size of the recombinant UGT11 protein at 57 kDa. Later, selected IMAC fractions were combined for SEC. D. SEC chromatogram. The x-axes show volume (mL) and SEC fractions. E. SDS-PAGE of SEC fractions 3 to 16. F. Western blot (WB) of SEC with anti-His-HRP antibody fractions 3 to 16. G. SDS-PAGE of SEC fractions 17 to 30. H. WB of SEC with anti-His-HRP antibody fractions 17 to 30.

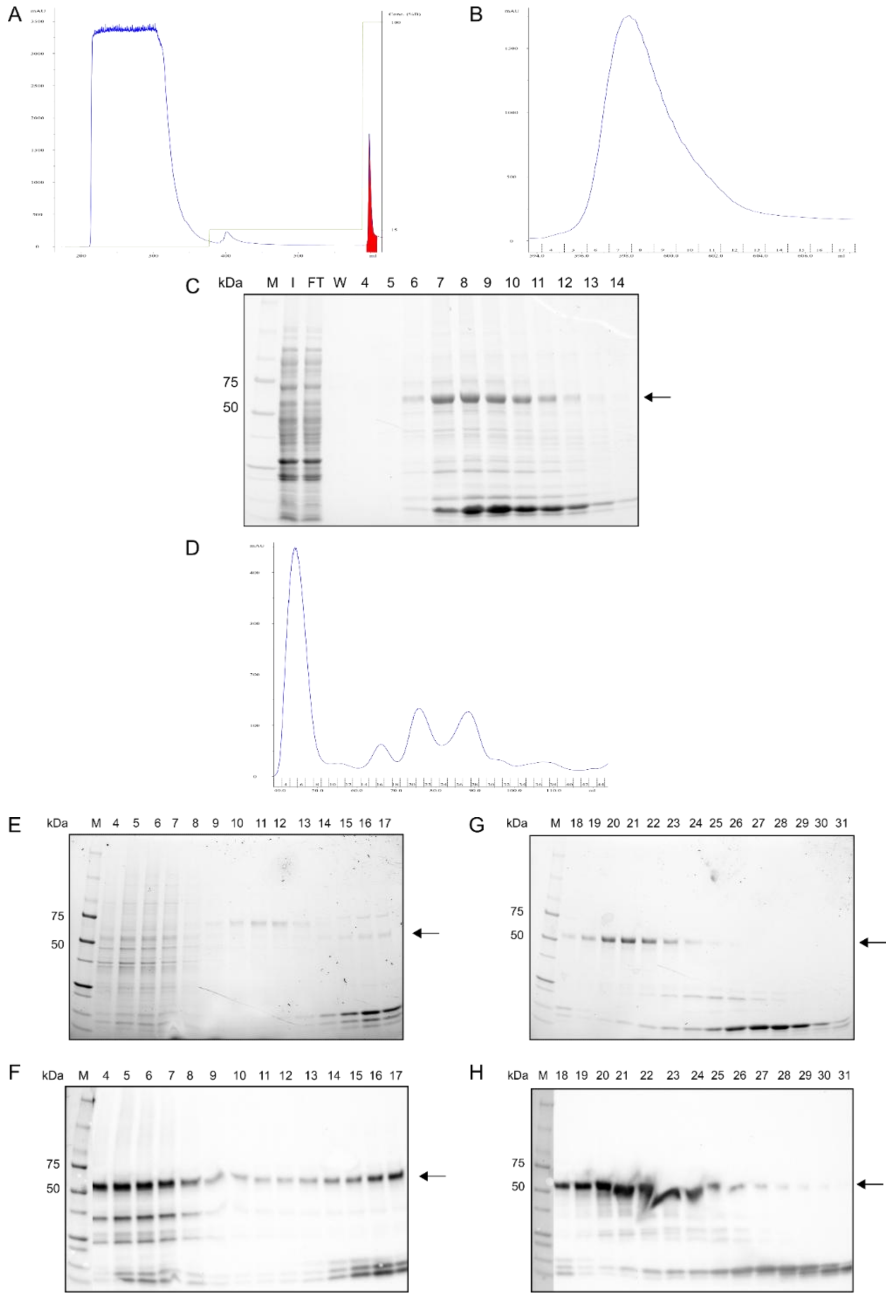


**Supplementary Fig. 8. Protein purification of SlUGT12.** SlUGT12 was fused to a His-tag, heterologously expressed in *E. coli* Rosetta (DE3), and purified by a two-step protein purification strategy of immobilized metal affinity chromatography (IMAC) and size exclusion chromatography (SEC). A. IMAC chromatogram. The chromatogram illustrates the absorption at 280 nm during elution (milli absorption, units mAU). The second y-axis shows elution buffer concentration (%). The fraction that contains the protein of interest is marked in red. B. IMAC chromatogram showing a zoom-in view of the peak containing the protein of interest. The x-axes show volume (mL) and IMAC fractions. The IMAC fractions were later used for sodium dodecyl sulfate (SDS)-polyacrylamide gel electrophoresis (PAGE). C. SDS-PAGE of IMAC fractions with samples of input (I), flow-through (FT), wash (W), and fractions 4 to 14. The black arrow highlights the size of the recombinant UGT12 protein at 58.7 kDa. Later, selected IMAC fractions were combined for SEC. D. SEC chromatogram. The x-axes show volume (mL) and SEC fractions. E. SDS-PAGE of SEC fractions 4 to 17. F. Western blot (WB) of SEC with anti-His-HRP antibody fractions 4 to 17. G. SDS-PAGE of SEC fractions 18 to 31. H. WB of SEC with anti-His-HRP antibody fractions 18 to 31.

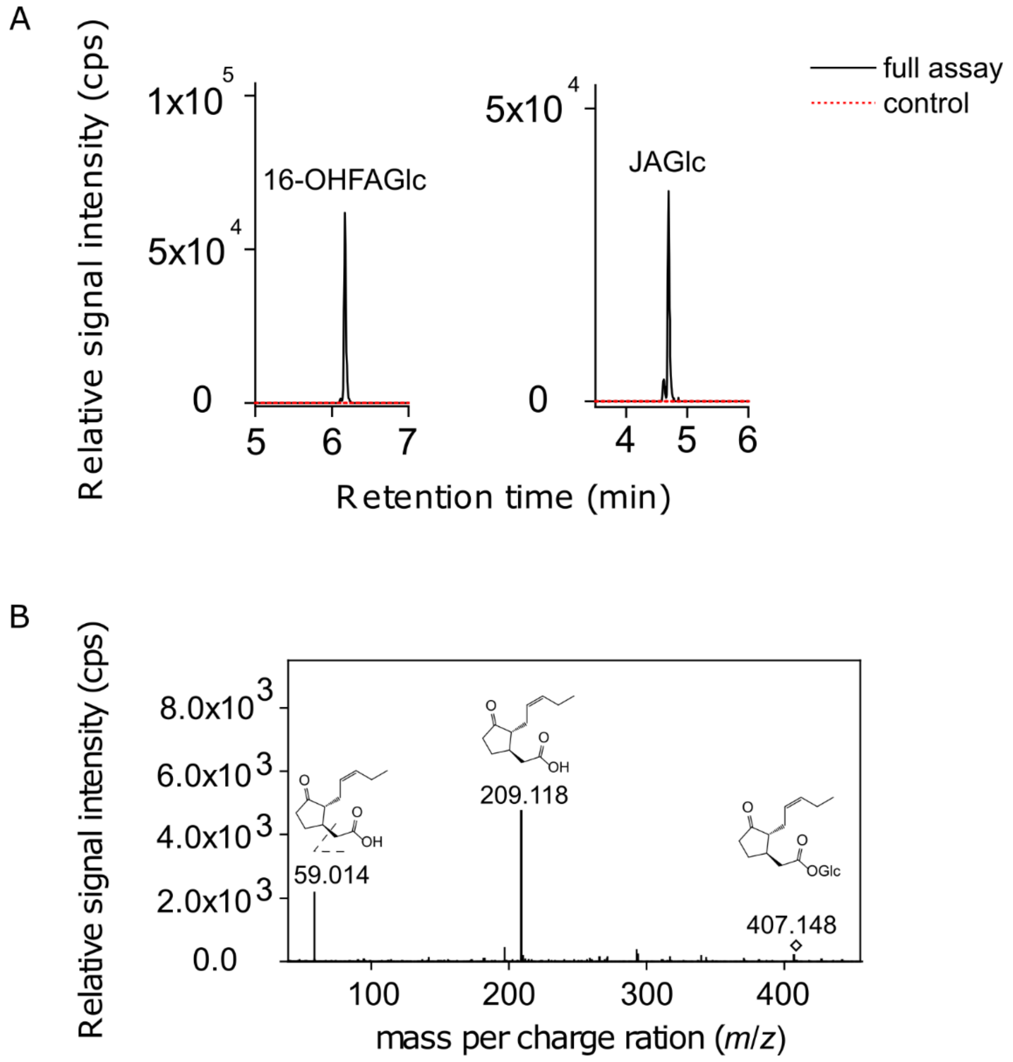


**Supplementary Fig. 9. LC-MS-based activity assays of SlUGT8 with JA and 12-OH-JA after 30 min of incubation.** A. JA activity assay using 0.5 mM JA and 5 mM UDP-Glc with 5 µg of purified SlUGT8 revealing production of JA-Glc. SlUGT10, SlUGT11, and SlUGT12 did not produce JA-Glc using the same experimental conditions and were not included in the figure. 16-OH-FA was used as activity control. B. JA-Glc parental and fragmented ions resulting from activity assays. The 30-min reactions were stopped by adding 50 µL of MeOH and analyzed by LC-MS. Extracted ion chromatograms are shown for the product JA-Glc. Signal intensity is given in counts per second (cps).

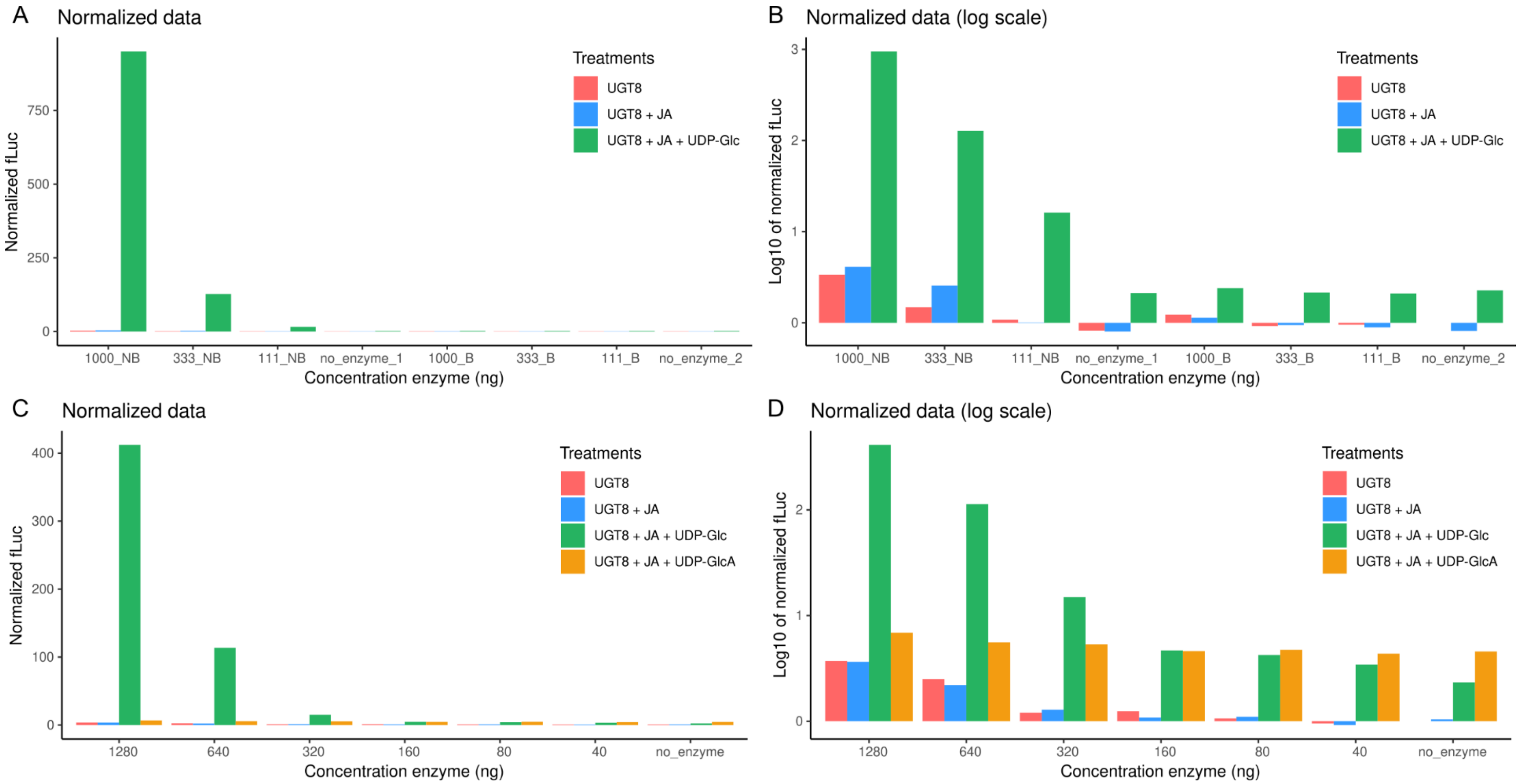


**Supplementary Figure 10. Luminescence-based activity assays of SlUGT8 with JA.** A. Activity assays to determine the minimal concentration of SlUGT8 using JA and UDP-Glc as substrates and three concentrations of SlUGT8 (denoted as non-boiled protein, NB), and negative control (denoted as boiled protein, B). B. Same data as in A, but data is presented in a log scale. C. Activity assays to determine substrate specificity using JA and two sugars, UDP-glucose (UPD-Glc) or UDP-glucuronic acid (UDP-GlcA), as substrates. Three concentrations of SlUGT8 (NB), and negative control (B) were used. D. Same data as in C but presented in log scale. In each experiment, samples incubated with only SlUGT8, and incubated with only SlUGT8 and JA were included as controls. For these preliminary experiments, only one repetition for each treatment was performed and therefore no statistical analyses were executed.

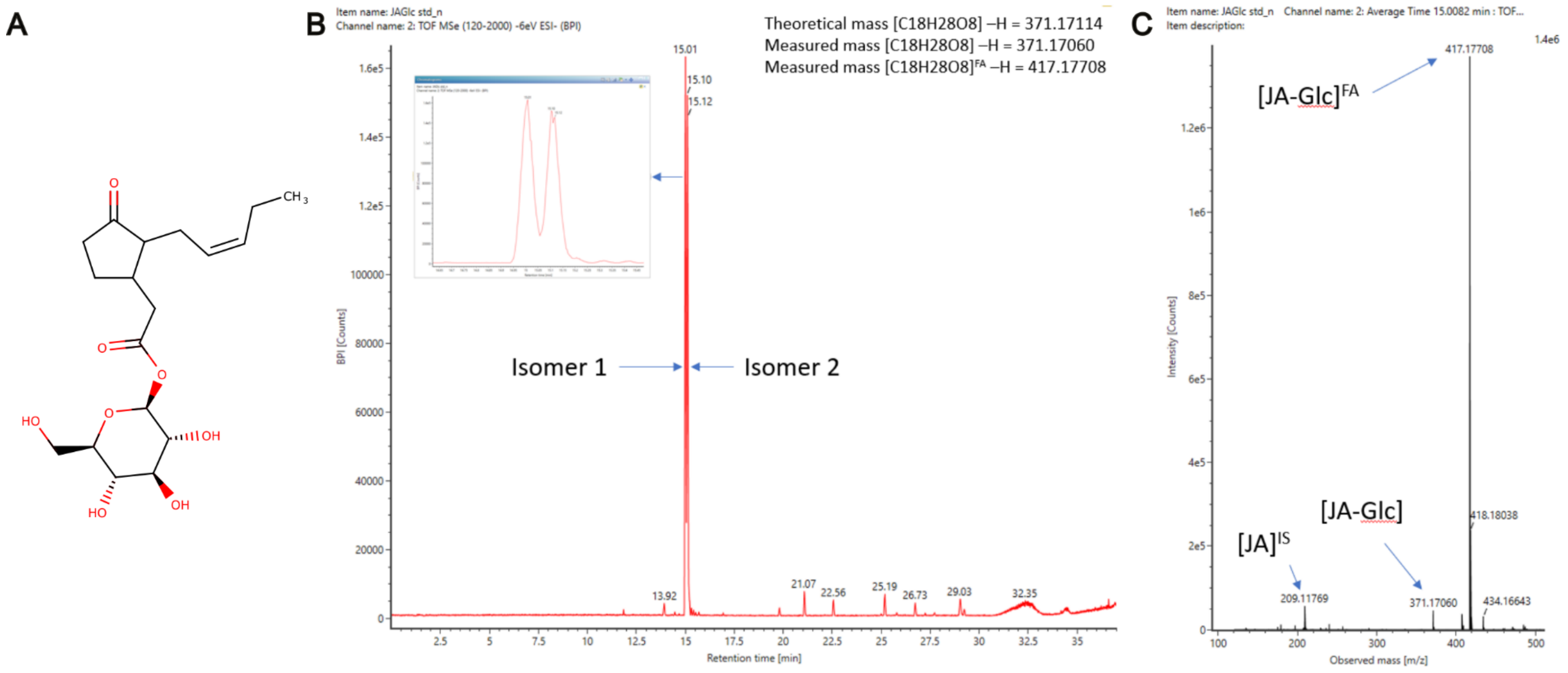


**Supplementary Fig. 11. Chemical synthesis and detection of jasmonyl-1-β-glucose (JA-Glc).** A. Chemical structure of the synthesized JA-Glc. B. LC-MS extracted chromatogram for the product of the chemical JA-Glc synthesis, depicting two isomers and the product of the commercial racemic JA used. C. LC-MS extracted chromatogram showing the mass measured of the product of the JA-Glc synthesis displaying the formic acid (FA) adduct of the JA-Glc as the highest peak and other fragments generated. IS: internal standard.

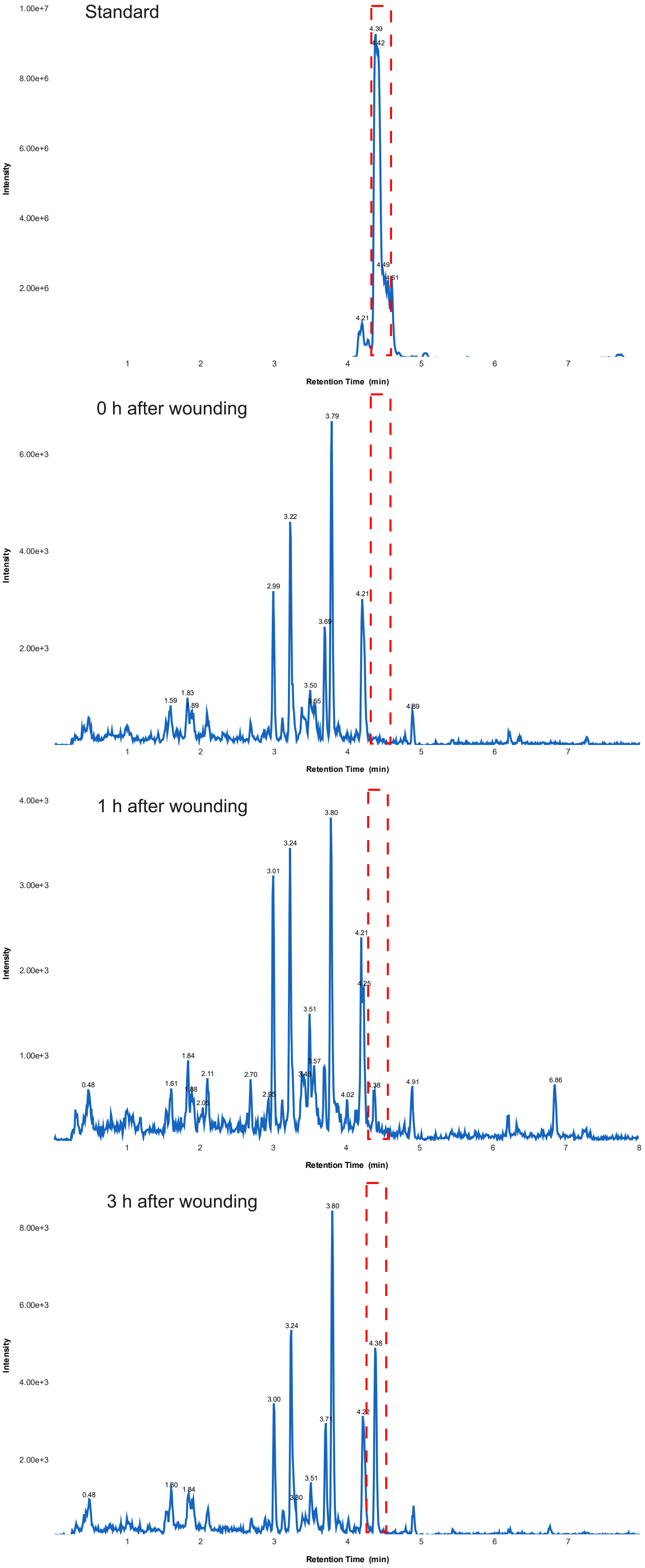
Standard
0 h after wounding
1 h after wounding
3 h after wounding
Intensity
Retention Time (min)
1.00e+7
8.00e+6
6.00e+6
4.00e+6
2.00e+6
4.39
4.42
4.49
4.61
4.21
6.00e+3
4.00e+3
2.00e+3
3.79
3.22
2.99
4.21
3.69
3.50
3.55
1.83
1.59
1.89
4.89
3.00e+3
1.00e+3
3.80
3.24
3.01
3.51
3.57
1.84
0.48
1.61
2.11
2.70
4.38
4.91
6.86
4.02
8.00e+3
4.38
3.77
4.22
3.00
1.60
3.51

**Supplementary Fig. 12. JA-Glc analysis of wounded tomato leaves**. The four extracted ion chromatograms from a TOF-MRM experiment with the transition *m/z* 417.17 → *m/z* 209.12 with a collision energy of 30 eV. The top chromatogram represents the JA-Glc standard, and the three chromatograms below it are for extracts from a single leaf (biological replicate) at 0 h, 1 h, and 3 h after wounding. The peak for jasmonoyl-β-D-Glc (JA-Glc) is marked with a red dashed-line box ($t_R$ = 4.38 - 4.39 min).

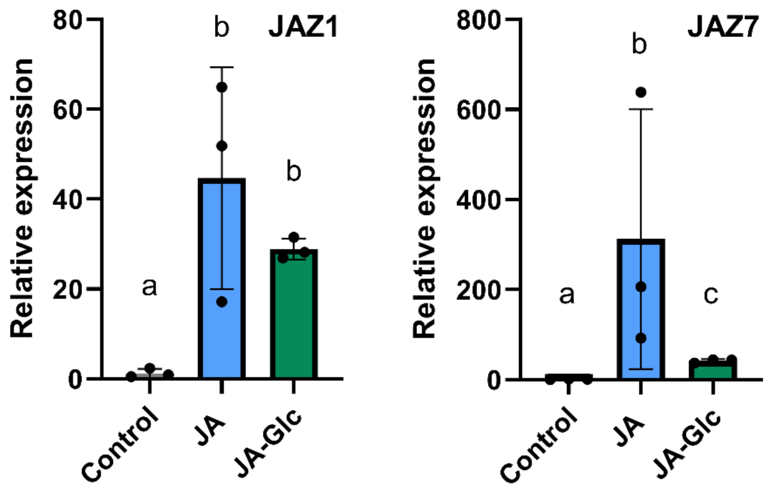


**Supplementary Fig. 13. qRT-PCR analysis of the JA-responsive marker genes in tomato roots after treatment with JA and JA-Glc.** Three wild-type roots were employed for each 6-h treatment with 1 µM JA, 1 µM JA-Glc, or ethanol as control. Black dots represent values for each biological replicate, bars represent means, and error bars display standard deviation. Lower case letters represent statistical differences between treatments calculated by Tukey's HSD test, following one-way ANOVA (p-value < 0.05).

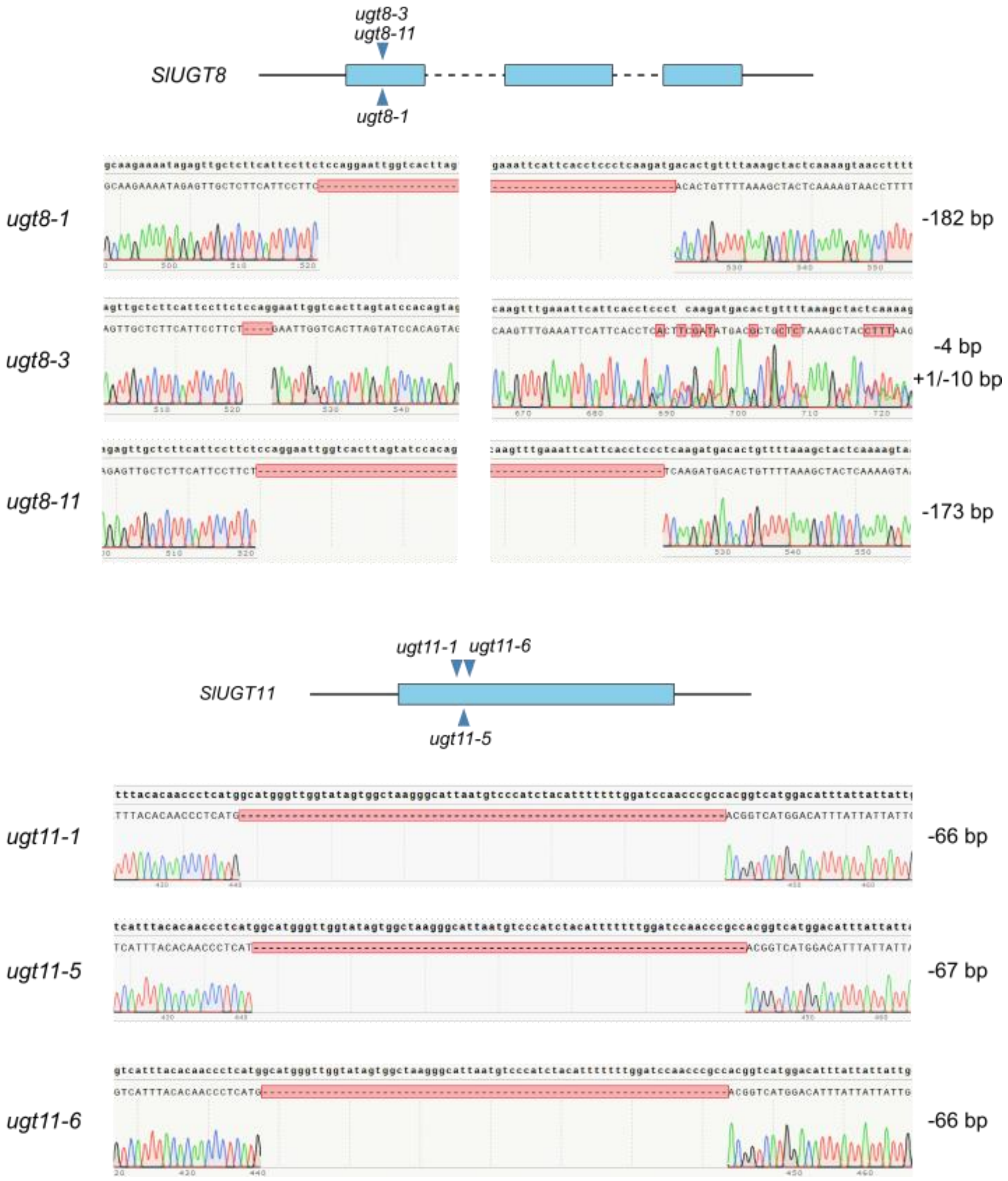


**Supplementary Fig. 14. Overview of mutations of tomato hairy root CRISPR-generated mutants**. Blue squares show coding sequence exons of *SlUGT* genes targeted by CRISPR. Blue arrow heads point to the positions of mutations, and the results of Sanger sequencing for each mutant line are shown below.

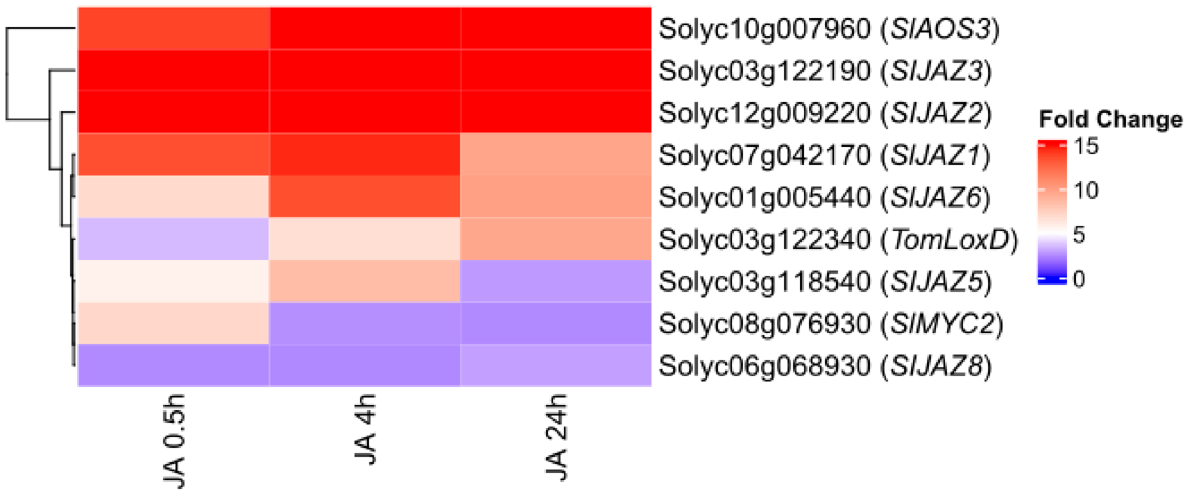


**Supplementary Fig. 15. Heatmap showing gene expression patterns of four known JA biosynthetic and signaling genes in an RNA-Seq experiment following JA treatment at three time points.** The heatmap depicts the fold-change of JA-treated samples compared to mock at 0.5, 4, and 24 h after treatment. Upregulation is depicted in white and red (fold-change > 2 and FDR < 0.05), and no effect in blue (n = 3).

**Supplementary Table 1. Differentially abundant peptides of the protein A0A3Q7EA02 (Solyc01g009990) upon LiP-MS with 10 mM JA.** A0A3Q7EA02 was detected as the first tomato homolog of the Arabidopsis cyclophilin 20-3 (CYP20-3, At3g62030).

| Peptide sequence | UniProt ID | Protein name | Adjusted p-value | Log2 fold-change | Solyc ID |
|---|---|---|---|---|---|
| AVEVADLQSK | A0A3Q7EA02 | Peptidyl-prolyl cis-trans isomerase (PPIase) (EC 5.2.1.8) | 0.00005 | 7.54 | Solyc01g009990 |
| LVHTGPGIVSM | A0A3Q7EA02 | Peptidyl-prolyl cis-trans isomerase (PPIase) (EC 5.2.1.8) | 0.00025 | 4.42 | Solyc01g009990 |
| DFMIQGGDFDKGNGTGGK | A0A3Q7EA02 | Peptidyl-prolyl cis-trans isomerase (PPIase) (EC 5.2.1.8) | 0.00156 | -1.95 | Solyc01g009990 |

**Supplementary Table 2. Differentially abundant peptides of proteins known to be involved in JA biosynthesis upon LiP-MS with 10 mM JA.**

| Peptide sequence | UniProt ID | Protein name | Adjusted p-value | Log2 fold-change | Solyc ID |
|---|---|---|---|---|---|
| MLGQLVGGLIGGHHDSK | P38415 | Linoleate 9S-lipoxygenase A (EC 1.13.11.58) (SlLOX) | 0.0006784 | -2.9 | Solyc08g014000 |
| LTTDEIPQIVNEFR | Q9XG54 | 12-oxophytodienoate reductase 1 (EC 1.3.1.42) (12-oxophytodienoate-10,11-reductase 1) (OPDA-reductase 1) (SlOPR1) | 0.0004082 | -3.78 | Solyc10g086220 |
| YLTAIKDPDQR | A0A3Q7IB30 | acyl-CoA oxidase activity (SlACX) | 0.0059306 | 2.45 | Solyc10g008110 |
| MAAFYAGFPDTVPIR | A0A3Q7I392 | 3-ketoacyl CoA thiolase (SlKAT ) | 0.0049945 | -1.25 | Solyc09g061840 |
| ATSLPLVPPILVALVNNADWIK | A0A3Q7JD55 | OPC-8:0-CoA ligase (SlOPCL) | 0.0055531 | -2.3 | Solyc12g094520 |

**Supplementary Table 3. List of all identified peptides from SlUGT8, SlUGT9, SlUGT10, SlUGT11, and SlUGT12 in the LiP-MS experiment with 10 mM JA.**

| UGT | Sequence | UniProt_ID | SolGenomics_ID | Adjusted p-value | Log2_fold-change |
|---|---|---|---|---|---|
| SlUGT8 | VIGWAPQLAILSHK* | A0A3Q7J5C5 | Solyc12g014010 | 0.0037 | -2.14 |
| SlUGT8 | IELLFIPSPGIGHLVSTVEMAK* | A0A3Q7J5C5 | Solyc12g014010 | 0.0435 | -1.46 |
| SlUGT8 | DATLPGDYENFEEVLPEGFLQR | A0A3Q7J5C5 | Solyc12g014010 | 0.1700 | -1.31 |
| SlUGT8 | FIHLPQDDTVLK | A0A3Q7J5C5 | Solyc12g014010 | 0.2411 | -1.32 |
| SlUGT9 | DSANDFNILGFSNPVPIK* | A0A3Q7F5R7 | Solyc02g081690 | 0.0081 | -1.98 |
| SlUGT9 | AGHPFLWAIR* | A0A3Q7F5R7 | Solyc02g081690 | 0.0360 | -4.20 |
| SlUGT9 | ELPNDYTNLEEILPNGFLESTK | A0A3Q7F5R7 | Solyc02g081690 | 0.0540 | -0.90 |
| SlUGT9 | WLDNQEPSSVIFLSFGSMGSLK | A0A3Q7F5R7 | Solyc02g081690 | 0.0745 | -2.22 |
| SlUGT9 | HIDGYSWFLHHAK | A0A3Q7F5R7 | Solyc02g081690 | 0.2737 | -1.00 |
| SlUGT9 | RELPNDYTNLEEILPN | A0A3Q7F5R7 | Solyc02g081690 | 0.5196 | -0.77 |
| SlUGT9 | VVFISTPALGNIVPIIEFAK | A0A3Q7F5R7 | Solyc02g081690 | 0.7258 | -0.89 |
| SlUGT9 | RELPNDYTNLEEILPNGFLESTK | A0A3Q7F5R7 | Solyc02g081690 | 0.9315 | 0.12 |
| SlUGT10 | VNEENGIVGR* | D7S016 | Solyc01g095620 | 0.0020 | 4.26 |
| SlUGT10 | FFNVQDSTNPLEFLPK* | D7S016 | Solyc01g095620 | 0.0176 | -1.29 |
| SlUGT10 | IAILPSPGMGHLIPLVEFAK* | D7S016 | Solyc01g095620 | 0.0182 | -2.28 |
| SlUGT10 | GFGLVLPNWAPQAR | D7S016 | Solyc01g095620 | 0.1385 | -1.35 |
| SlUGT10 | IANATFFNVQDSTNPLEFLPK | D7S016 | Solyc01g095620 | 0.1531 | -0.92 |
| SlUGT10 | AQIPHIAILPSPGMGHLIPLVEFAK | D7S016 | Solyc01g095620 | 0.1536 | -1.94 |
| SlUGT10 | NVQDSTNPLEFLPK | D7S016 | Solyc01g095620 | 0.1617 | 2.22 |
| SlUGT10 | DSTNPLEFLPK | D7S016 | Solyc01g095620 | 0.2424 | -1.56 |
| SlUGT10 | ELEGGAIGALQK | D7S016 | Solyc01g095620 | 0.6871 | 0.24 |
| SlUGT11 | SIIDQIELLNSEENPR* | A0A3Q7IA70 | Solyc09g092480 | 0.0035 | -1.11 |
| SlUGT11 | VLVNTFDDLEFDALR* | A0A3Q7IA70 | Solyc09g092480 | 0.0101 | -0.83 |
| SlUGT11 | DFPSFVFTDVNSK* | A0A3Q7IA70 | Solyc09g092480 | 0.0136 | -1.09 |
| SlUGT11 | NLTMVGIGPSIPSAFLDGNDPLDK | A0A3Q7IA70 | Solyc09g092480 | 0.0908 | -1.56 |
| SlUGT11 | VNTFDDLEFDALR | A0A3Q7IA70 | Solyc09g092480 | 0.1611 | -0.40 |
| SlUGT11 | TFDDLEFDALR | A0A3Q7IA70 | Solyc09g092480 | 0.2686 | -1.46 |
| SlUGT11 | SFTPFSDGYDGK | A0A3Q7IA70 | Solyc09g092480 | 0.3925 | 0.26 |
| SlUGT11 | SIIDQIELL | A0A3Q7IA70 | Solyc09g092480 | 0.4327 | -0.99 |
| SlUGT11 | VGIGPSIPSAFLDGNDPLDK | A0A3Q7IA70 | Solyc09g092480 | 0.5281 | -0.57 |
| SlUGT12 | TTNPEEYMIK* | A0A3Q7JX81 | Solyc12g057060 | 0.0003 | 4.51 |
| SlUGT12 | VINDVLLSSK* | A0A3Q7JX81 | Solyc12g057060 | 0.0195 | 1.65 |

*Significant peptides with q-value <0.05. These significant peptides were highlighted in the metabolite–protein complex 3D structure models (Figure 3 and Supplemental Figures 2-4).

**Supplementary Table 4. Results of the Inference of CRISPR Edits (ICE) analysis (Synthego) of CRISPR/Cas9 lines targeting all five *SlUGTs*.** ICE indicates indel percentage, KO-Score is a percentage of frame-shift mutations or deletions of 21 bp or larger, $R^2$ is the correlation of indel distribution predicted by ICE and the Sanger sequencing data, mean discord scores demonstrate discord before and after the gRNA sites. Indels detected are indicated by the indel size shown with single quotation marks and the percentage of each indel after the colon.

| Line | Gene | ICE | KO-Score | $R^2$ | Mean Discord Before | Mean Discord After | Guide Sequences | Control Sample Quality Score | Edit Sample Quality Score | Indels |
|---|---|---|---|---|---|---|---|---|---|---|
| 5 | *SlUGT8* | 100 | 100 | 0.4 | 0.018146503 | 0.748128314 | CCCTCAAGATGACACTGTTT,TCTCCAGGAATTGGTCACTT | 61 | 61 | '1': 13.0, '-2': 12.0, '-35': 2.0, '-172': 64.0, '-188': 3.0, '-192': 1.0, '-193': 2.0, '-198': 1.0, '-200': 2.0 |
| 5 | *SlUGT9* | 100 | 94 | 0.94 | 0.084311155 | 0.709249404 | CCTTATTTCAGACTCGGACT,GCGATGGTGGATCCGCCACG | 58 | 59 | '6': 6.0, '-17': 1.0, '-134': 93.0 |
| 5 | *SlUGT10* | 100 | 82 | 0.58 | 0.058232653 | 0.759170494 | CCGTGGATCGGAGTACAACC,CGACACCGTTTCATCAAGGT | 58 | 62 | '1': 18.0, '3': 1.0, '-1': 7.0, '-2': 6.0, '-3': 16.0, '12': 1.0, '17': 2.0, '-17': 2.0, '-60': 32.0, '-61': 8.0, '-63': 4.0, '-84': 2.0, '-87': 1.0 |
| 5 | *SlUGT11* | 100 | 100 | 0.4 | 0.016972945 | 0.725754248 | GCCACGGTCATGGACATTTA,CATGGCATGGGTTGGTATAG | 61 | 58 | '8': 3.0, '-2': 10.0, '-4': 26.0, '-22': 2.0, '-37': 6.0, '-71': 52.0, '-83': 1.0 |
| 5 | *SlUGT12* | 89 | 64 | 0.71 | 0.019383556 | 0.504672116 | TATTCCACTGTTATCGAAAA,TGTAAAGCTCATAACACCAT | 59 | 62 | '0': 11.0, '-1': 33.0, '-3': 14.0, '-5': 18.0, '-6': 11.0, '13': 3.0, '-11': 9.0, '-31': 1.0 |
| 6 | *SlUGT8* | 100 | 100 | 0.51 | 0.011796922 | 0.717970896 | CCCTCAAGATGACACTGTTT,TCTCCAGGAATTGGTCACTT | 61 | 61 | '1': 18.0, '-10': 32.0, '-21': 44.0, '-23': 1.0, '-35': 1.0, '-188': 3.0, '-206': 1.0 |

| | | | | | | | | | | |
|---|---|---|---|---|---|---|---|---|---|---|
| 6 | *SlUGT9* | 100 | 100 | 0.79 | 0.058656783 | 0.729728408 | CCTTATTTCAGACTCGGACT,GCGATGGTGGATCCGCCACG | 58 | 37 | '2': 100.0 |
| 6 | *SlUGT10* | 0 | 0 | 1 | 0.112567893 | 0.031574643 | CCGTGGATCGGAGTACAACC,CGACACCGTTTCATCAAGGT | 58 | 23 | '0': 100.0, '-1': 0.0 |
| 6 | *SlUGT11* | 53 | 53 | 0.86 | 0.012609972 | 0.37616677 | GCCACGGTCATGGACATTTA,CATGGCATGGGTTGGTATAG | 61 | 61 | '0': 47.0, '-1': 53.0 |
| 6 | *SlUGT12* | 0 | 0 | 1 | 0.003682674 | 0.008004036 | TATTCCACTGTTATCGAAAA,TGTAAAGCTCATAACACCAT | 59 | 58 | '0': 100.0, '-147': 0.0 |


1 Wu, Q., Peng, Z., Zhang, Y. & Yang, J. COACH-D: improved protein-ligand binding sites prediction with refined ligand-binding poses through molecular docking. *Nucleic Acids Res* **46**, W438–W442 (2018). https://doi.org/10.1093/nar/gky439
